# A triple distinction of cerebellar function for oculomotor learning and fatigue compensation

**DOI:** 10.1101/2022.06.15.496290

**Authors:** Jana Masselink, Alexis Cheviet, Denis Pélisson, Markus Lappe

## Abstract

The cerebellum implements error-based motor learning via synaptic gain adaptation of an inverse model, i.e. the mapping of a spatial movement goal onto a motor command. Recently, we modeled the motor and perceptual changes during learning of saccadic eye movements, showing that learning is actually a threefold process. Besides motor recalibration of the inverse model (1), learning also comprises perceptual recalibration of the visuospatial target map (2) and of a forward dynamics model that estimates the saccade size from corollary discharge (3). Yet, the site of perceptual recalibration remains unclear. Here we dissociate cerebellar contributions to the three stages of learning by modeling the learning data of eight cerebellar patients and eight healthy controls. Results showed that cerebellar pathology restrains short-term recalibration of the visuospatial target map and of the inverse model while the forward dynamics model is well informed about the reduced saccade change. Moreover, patients showed uncompensated oculomotor fatigue caused by insufficient upregulation of saccade duration. According to our model, this could induce long-term perceptual compensation, consistent with the overestimation of target eccentricity found in the patients’ baseline data. We conclude that the cerebellum mediates short-term adaptation of the visuospatial target map and of the inverse model, especially by control of saccade duration. The forward dynamics model was not affected by cerebellar pathology.

**Author Summary:** Achieving a fine-grained understanding of how the cerebellum continuously recalibrates our movements is an ongoing challenge in sensorimotor neuroscience. Recently, we showed that recalibration of saccadic eye movements does not only operate in motor space, i.e. by adjusting the motor command, but also in external and internal visual space, i.e. by adjusting the spatial representation of the target and the internal saccade size. For this purpose, (1) we developed a paradigm that allowed us to monitor changes of the internal saccade size estimated from trans-saccadic target localizations, and (2) we unified the three learning processes in one computational modeling framework. Here we apply this approach to the saccade learning data of patients with a neurodegenerative cerebellar disease. First, we dissociate the cerebellar role in recalibration of these three sites of learning. Second, we show how learning is transposed to saccade kinematics. Third, we provide first insights into the perceptual consequences of cerebellar pathology that, according to our model, may be a mechanism to recover from disease-specific motor deficits. Our modeling framework may help to dissociate the contribution of specific sensorimotor areas to adaptive behavior as well as to improve the understanding of learning deficits and compensatory strategies in the clinical context.

## Introduction

The cerebellum modulates supervised error-based learning (Marr, 1969; Albus, 1971; Herzfeld, Kojima, Soetedjo, & Shadmehr, 2015). If a movement is inaccurate, an error signal adapts the synaptic gains of cerebellar Purkinje cells (Bays & Wolpert, 2007; Herzfeld, Kojima, Soetedjo, & Shadmehr, 2018; Raymond & Medina, 2018; Markanday, Inoue, Dicke, & Thier, 2021) that implement an inverse model, i.e. the mapping of the visuospatial movement goal onto a motor command (Kawato, 1999; Bays & Wolpert, 2007; Franklin & Wolpert, 2011). If e.g. the eye muscles fatigue, the synaptic gains are modulated such that the saccade does not fall short of the target (McLaughlin, 1967; Havermann & Lappe, 2010; Pelisson, Alahyane, Panouilleres, & Tilikete, 2010). To compensate for continuous fluctuations of muscle dynamics, the cerebellum is constantly recalibrating the saccadic circuitry (Srimal, Diedrichsen, Ryklin, & Curtis, 2008; Shadmehr, Smith, & Krakauer, 2010; Wolpert, Diedrichsen, & Flanagan, 2011).

However, the traditional view on the cerebellum as a pure motor hub has changed. The cerebellum is interconnected with almost the entire cortex (Middleton & Strick, 2001; Hoshi, Tremblay, Féger, Carras, & Strick, 2005; Bostan, Dum, & Strick, 2010), including retinotopic areas (Buckner, Krienen, Castellanos, Diaz, & Yeo, 2011; Brissenden, Levin, Osher, Halko, & Somers, 2016; Marek et al., 2018), and basal ganglia (?, ?), and holds at least five visuospatial maps itself (van Es, van der Zwaag, & Knapen, 2019). Moreover, it has been shown that saccadic motor learning also comprises changes in visuospatial representations. These changes occur, firstly, in the visuospatial target map when errors are assigned to an internal failure of spatial target representation (Moidell & Bedell, 1988; Collins, Dore-Mazars, & Lappe, 2007; Schnier, Zimmermann, & Lappe, 2010; Zimmermann & Lappe, 2010; Gremmler, Bosco, Fattori, & Lappe, 2014). Secondly, these changes occur in the internal representation of saccade size as measured by trans-saccadic target localizations (Bahcall & Kowler, 1999; Zimmermann & Lappe, 2009; Schnier et al., 2010).

Changes in the internal saccade representation affect the forward dynamics model that transforms the corollary discharge, i.e. a copy of the motor command (*CD_M_*; Duhamel, Colby, & Goldberg, 1992; Umeno & Goldberg, 1997; Crapse & Sommer, 2009) into a visuospatial representation of saccade size (*CD_V_*; Bays & Wolpert, 2007; Sommer & Wurtz, 2008; Crapse & Sommer, 2008). Intact *CD_V_* information is necessary to correctly bridge the spatial gap between pre- and post-saccadic visual input that supports our perception of a stable world (Sperry, 1950; von Holst & Mittelstaedt, 1950; Wurtz, 2008). In Masselink and Lappe (2021), we implemented the action-perception entanglement during learning in a computational model that collectively adapts the visuospatial target map, the inverse model and the forward dynamics model in response to postdictive motor error, i.e. motor error with respect to a postdicted target position.

Cerebellar perturbations restrain saccade changes during learning, as shown e.g. in lesioned monkeys (Takagi, Zee, & Tamargo, 1998; Barash et al., 1999), in cerebellar patients (Alahyane et al., 2008; Golla et al., 2008; Xu-Wilson, Chen-Harris, Zee, & Shadmehr, 2009) and in healthy subjects with non-invasive cerebellar stimulation (Panouilleres et al., 2012; Avila et al., 2015; Jenkinson & Miall, 2010). Because of its circular learning architecture, its visuospatial maps and its CD projection pathway to frontal cortex (Middleton & Strick, 2000), the cerebellum is an ideal candidate to also recalibrate the visuospatial target map and the forward dynamics model. In Cheviet et al. (2022), we found first evidence that changes in visuospatial representations during saccadic motor learning are impaired in cerebellar patients.

Here we use an expanded version of the visuomotor learning model of Masselink and Lappe (2021) to quantify the recalibration of the visuospatial target map, of the inverse model and of the forward dynamics model of eight patients with a neurodegenerative cerebellar disease. Data were taken from Cheviet et al. (2022), including eight healthy control subjects. To achieve a deeper understanding of which control processes are perturbed, we expanded the model of Masselink and Lappe (2021) with (1) how changes in the motor command are transposed to saccade kinematics (i.e. saccade peak velocity and saccade duration), and (2) how the visuomotor system compensates for a fatigue-induced decline in saccade peak velocity. Conditions comprise (1) learning from peri-saccadic inward and (2) from peri-saccadic outward target steps, and (3) compensation of neural oculomotor fatigue, i.e. without target step. Our results reveal that cerebellar pathology reduces short-term learning of the visuospatial target map and of the inverse model, especially when controlled by upregulation of saccade duration. Notably, the forward dynamics model and thus, the *CD_V_* signal, seemed correctly informed about the reduced saccade change. Moreover, we show that cerebellar pathology is accompanied by an overestimation of target eccentricity that, according to our model, could stem from error reduction to counteract oculomotor fatigue on a long timescale. We conclude that intact recalibration of the visuospatial target map and of the inverse model, particularly via control of saccade duration, depend on cerebellar integrity. The forward dynamics model, at least for learning from post-saccadic errors, is not affected by cerebellar pathology. Long-term learning at perceptual level may partially compensate for cerebellar motor deficits.

## Results

To reveal how adaptation of the visuospatial target map, of the inverse model and of the forward dynamics model for saccade motor learning relies on cerebellar integrity, we fitted a visuomotor learning model to the saccade and visual target localization data of eight cerebellar patients and eight healthy control subjects.

### Experimental data

Each subject participated in three conditions requiring saccades to a 20° rightward target as well as pre- and post-saccadic visual localizations of a shortly flashed bar presented around the usual target position (Figure 1A). In the pre-saccadic localization trials, subjects localized the flash with a mouse cursor while holding gaze at the fixation point. In the post-saccadic localization trials, subjects performed a saccade to a 20° rightward target and then localized the pre-saccadic flash that had appeared before target onset. The pre- and the post-exposure phase (phase 1 and 3) measured subjects’ state of saccade vector, pre- and post-saccadic visual localizations before and after exposure. The exposure phase (phase 2) induced either saccade shortening in response to (1) a 6° peri-saccadic inward target step (inward condition), (2) saccade lengthening in response to a 6° peri-saccadic outward target step (outward condition), or (3) tested the ability to maintain the saccade vector during repetitive, stereotyped saccades to a non-stepping target (no step condition). This task is known to induce oculomotor fatigue, i.e. a decline in saccade peak velocity. In healthy subjects, this is usually counteracted by upregulation of saccade duration such that the saccade vector stays stable (Schmidt, Abel, DellOsso, & Daroff, 1979; Fuchs & Binder, 1983; Bahill, Clark, & Stark, 1975; Straube, Robinson, & Fuchs, 1997; Chen-Harris, Joiner, Ethier, Zee, & Shadmehr, 2008; Xu-Wilson, Zee, & Shadmehr, 2009; Prsa, Dicke, & Thier, 2010). Compared to learning in response to an assumed fatigue of the eyes muscles, oculomotor fatigue is of neural origin and perhaps due to a loss of motivation or attention (Prsa et al., 2010). However, patients with cerebellar lesion of the vermis have shown to be impaired in compensating oculomotor fatigue (Golla et al., 2008).

**Figure 1.**
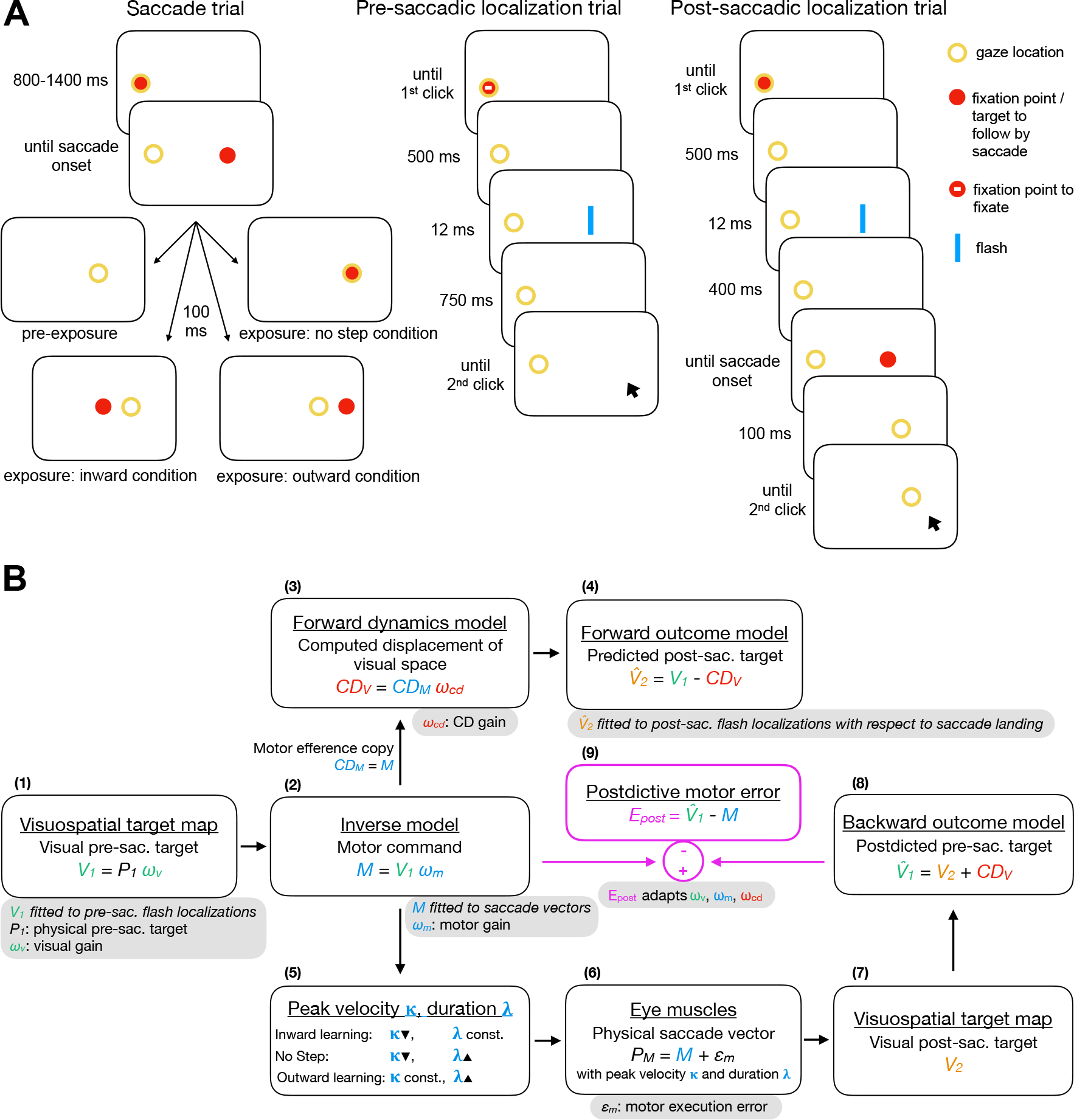
Experimental tasks and model framework. **(A)** In the saccade trials, subjects executed a saccade to a 20° rightward target. In the pre-exposure phase, the target was extinguished at saccade onset. In the exposure and post-exposure phase, the target either stepped 6° inward (inward condition), 6° outward (outward condition) or stayed at its initial position (no step condition). In the pre-saccadic localization trials, subjects localized a 12 ms flash with a mouse cursor while holding gaze at the fixation point. In the post-saccadic localization trials, subjects performed a saccade to a 20° rightward target and then localized the 12 ms pre-saccadic flash with a mouse cursor. Please note that the stimuli are not drawn to scale. The mouse cursor was a blue line pointer. The yellow circle illustrates gaze location but was not present at the stimulus display. **(B)** The target with the physical distance *P*_1_ is represented at the location *V*_1_ on the visuospatial map. An inverse model maps *V*_1_ onto a motor command *M*. Before saccade start, a forward dynamics model transforms a copy of the motor command *CD_M_* into visuospatial coordinates, i.e. into the computed displacement of visual space *CD_V_*. A forward outcome model then shifts retinal coordinates by *CD_V_* to predict the visual location of the post-saccadic target 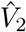. For saccade execution, the motor command is transposed to saccade peak velocity *κ* and saccade duration *λ*, producing the saccade vector *P_M_*. The actual post-saccadic target appears at retinal position *V*_2_. A backward outcome model then postdicts the visual post-saccadic target back to pre-saccadic space 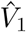. The visuomotor system evaluates the accuracy of the motor command with respect to the postdicted target position (*E_post_*) in order to adapt its gains *ω_v_*, *ω_m_* and *ω_cd_*. A decrease of *ω_m_* (inward learning) is controlled by downregulating saccade peak velocity while an increase of *ω_m_* (outward learning) is controlled by upregulation of saccade duration. When saccades are performed repetitively to the same, non-stepping target, oculomotor fatigue occurs, i.e. peak velocity declines, which is usually compensated by an increase of saccade duration.

Figure 2 shows the inward learning data of an example control subject (A) and of an example patient (B). Besides the higher variability of saccade endpoints (that has been reported before as a side-effect of cerebellar dysfunction, Barash et al., 1999; Golla et al., 2008; Xu-Wilson, Chen-Harris, et al., 2009; Thier & Markanday, 2019), the data show the impairment in amplitude adaptation of cerebellar patients compared to healthy controls.

**Figure 2.**
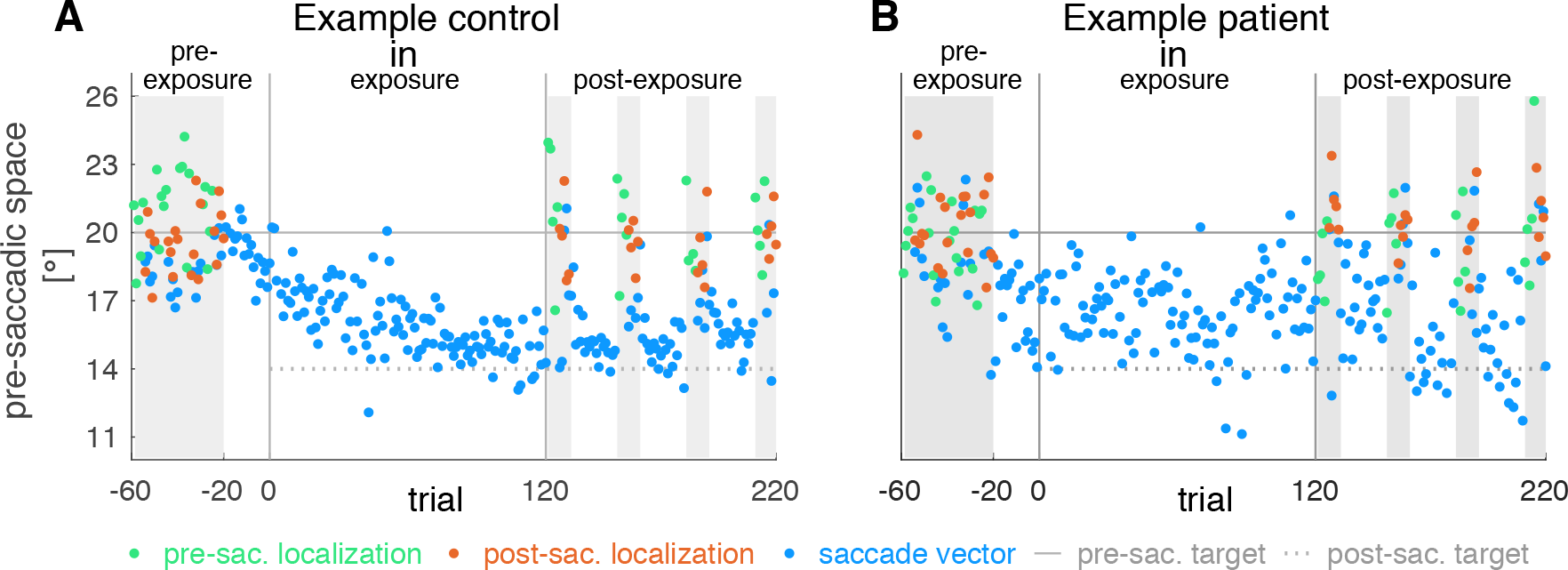
Example subject data for the inward condition. Saccade vectors, pre- and post-saccadic localizations for the inward condition of (**A**) an example control subject and (**B**) an example patient. The pre-exposure phase measured subjects’ baseline state without target step. The exposure phase induced saccadic learning and the post-exposure phase measured adaptation-induced changes to the saccade vector, pre- and post-saccadic localizations. Similar to the pattern that we found at group level, the patient shows less learning in the saccade vector and higher saccade endpoint variability compared to the control subject.

### Model fits

Figure 1B presents the framework of the expanded visuomotor learning model based on Masselink and Lappe (2021) that we fitted to the saccade vectors and pre- and postsaccadic visual localizations, separately for each group (controls, patients) and each condition (inward, no step, outward). The visual pre-saccadic target *V*_1_ (fitted to the pre-saccadic visual localizations) is represented on the visuospatial map with the visual gain *ω_v_* (Figure 1B-1). An inverse model transforms *V*_1_ into a motor command *M* (fitted to the saccade vectors) with the motor gain *ω_m_* (Figure 1B–2). Before saccade execution, a copy of the motor command is routed into the CD pathway where a forward dynamics model estimates the visuospatial size *CD_V_* of the upcoming saccade based on the CD gain *ω_cd_* (Figure 1B-3). A forward outcome model then shifts retinal coordinates by *CD_V_* to predict the retinal position of the post-saccadic target 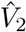 (fitted to the post-saccadic visual localizations; Figure 1B-4). For saccade execution, the motor command is transposed to saccade peak velocity *κ* and saccade duration *λ* (Figure 1B-5). After the saccade is performed (Figure 1B-6), the actual post-saccadic target appears at retinal position *V*_2_ (Figure 1B-7). A backward outcome model then postdicts the visual post-saccadic target back to pre-saccadic space 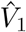 (Figure 1B-8). The postdictive motor error *E_post_* evaluates the accuracy of the motor command with respect to the postdicted target position in order to adapt its gains *ω_v_*, *ω_m_* and *ω_cd_* (Figure 1B-9). A decrease of *ω_m_* (inward learning) is controlled by downregulating saccade peak velocity while an increase of *ω_m_* (outward learning) is controlled by upregulation of saccade duration (Figure 1B-5).

The no step condition in which saccades are performed repetitively to the same, non-stepping target should lead to oculomotor fatigue, i.e. a decline in saccade peak velocity which is usually compensated by an increase of saccade duration. We captured peak velocity and duration changes in the no step condition by fitting a peak velocity decay rate *γ_κ_ ≥* 0 and a duration compensation rate *γ_λ_*. The latter describes the within-trial percentage by which saccade duration compensates for the peak velocity loss (0 *≤ γ_λ_ ≤* 1).

Figure 3 presents the model fits to the mean data of control subjects (A) and patients (B) for the three conditions (inward, no step, outward). Dots with errorbars show the mean data of the pre- and post-exposure phases, respectively, jagged blue lines show the trial-by-trial saccade data (A-1,B-1: saccade vectors; A-5,B-5: saccade peak velocities; A-6,B-6: saccade durations) and smooth lines show the trial-by-trial model fit to the data. Figure 4 depicts the baseline state as well as pre- to post-exposure changes for each condition, compared between patients and control subjects. Figure 5 shows the individual saccade peak velocities and Figure 6 the individual saccade durations in dependence on saccade vector for control subjects (C1-C8) and patients (P1-P8) from early trials (blue) to late trials (yellow). Table 1 gives an overview on the fitted parameters and residual standard errors as a measure of goodness of fit.

**Figure 3.**
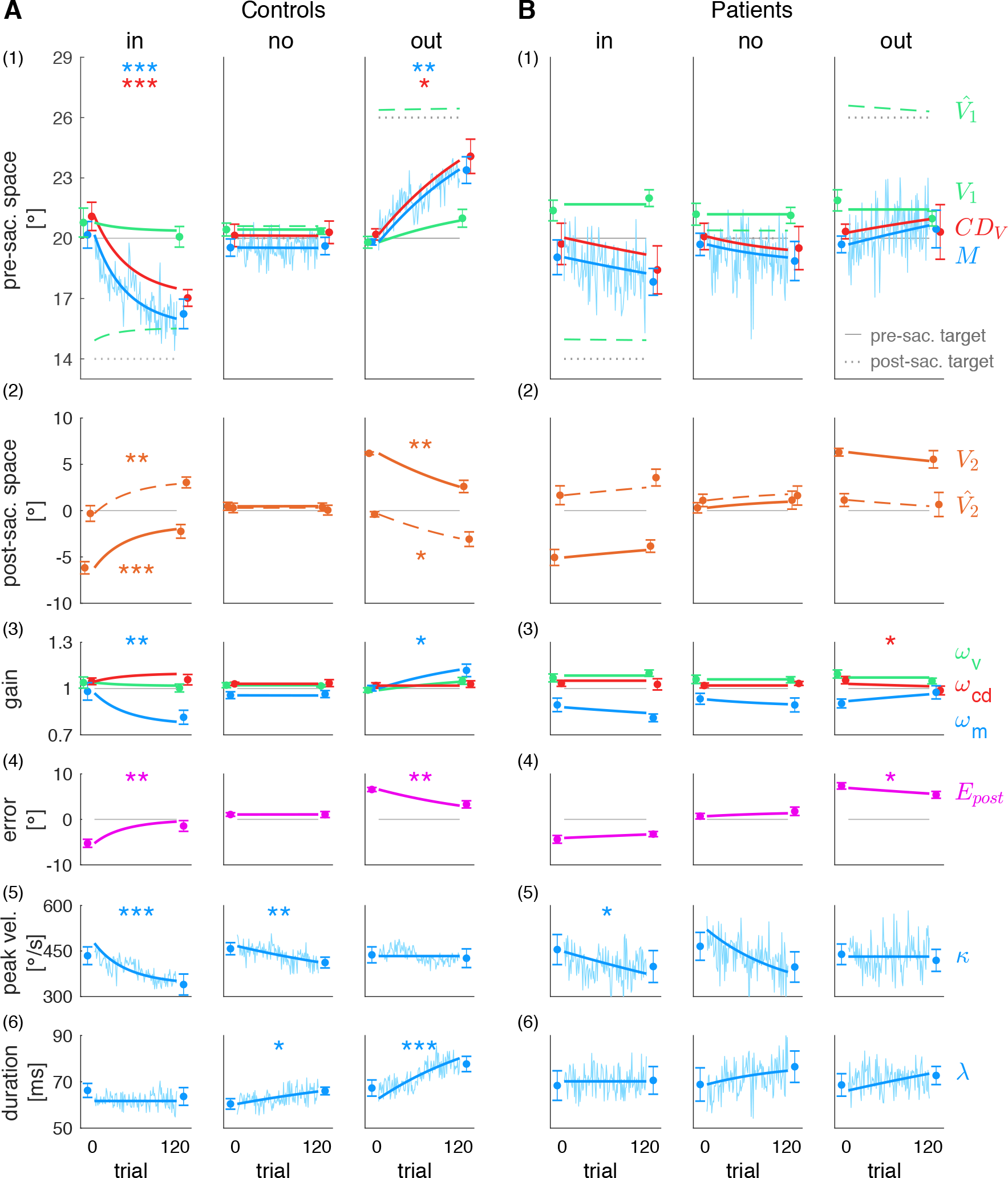
Model fits to the group data. Pre- and post-exposure mean standard error (dots with error bars), trial-by-trial saccade data of the exposure phase (jagged blue lines) and model fits to the data (smooth lines) for **(A)** control subjects and **(B)** cerebellar patients, separately for each condition (inward, no step, outward). (1) Visual pre-saccadic target *V*_1_ (fitted to pre-saccadic localizations), motor command *M* (fitted to saccade vectors), computed displacement of visual space *CD_V_*, postdicted pre-saccadic target 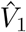. (2) Predicted post-saccadic target 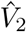 (fitted to post-saccadic localizations with respect to saccade landing), visual post-saccadic target *V*_2_. (3) Visual gain *ω_v_*, motor gain *ω_m_*, CD gain *ω_cd_*. (4) Postdictive motor error *E_post_*. (5) Saccade peak velocity *κ*. (6) Saccade duration *λ*. Asterisks indicate significant difference between the pre- and post-exposure phase with *** *p* < .001, ** *p* < .01, * *p* < .05 and n.s. *p ≥* .05.

**Figure 4.**
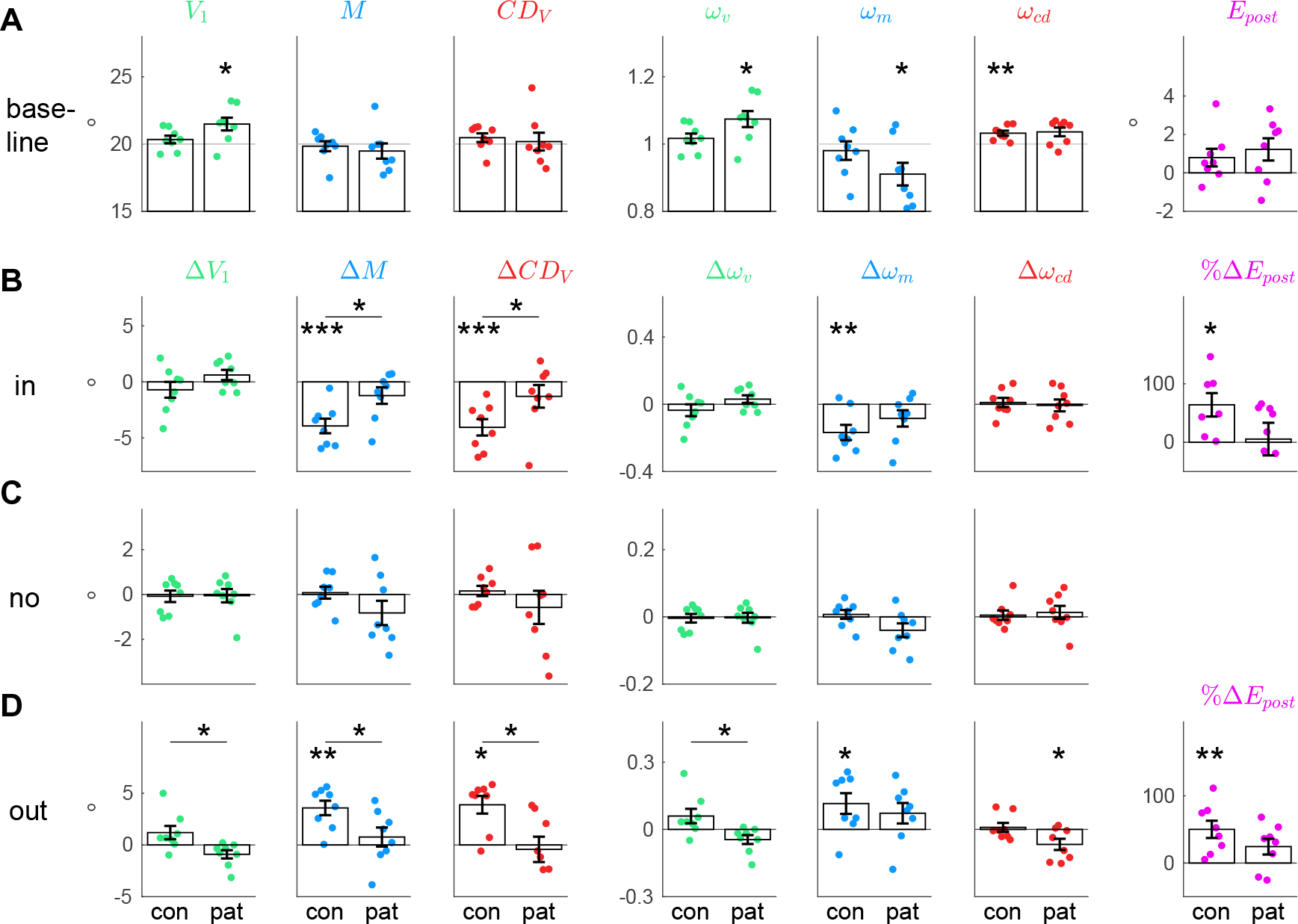
Baseline state and pre- to post-exposure changes. **(A)** Baseline state of the visuomotor system averaged across the pre-exposure phases of the three conditions (mean standard error), separately for control subjects and patients. Group-centered asterisks indicate significant difference from 20° (*V*_1_, *M*, *CD_V_*), from 1 (*ω_v_*, *ω_m_*, *ω_cd_*) or from zero (*E_post_*). **(B)** Pre- to post-exposure changes in the inward condition. **(C)** Pre- to post-exposure changes in the no step condition. **(D)** Pre- to post-exposure changes in the outward condition. For **B-D** group-centered asterisks indicate significant difference from zero. Panel-centered asterisks above a horizontal indicate significant difference between control subjects and patients with *** *p* < .001, ** *p* < .01, * *p* < .05 and n.s. *p ≥* .05.

**Figure 5.**
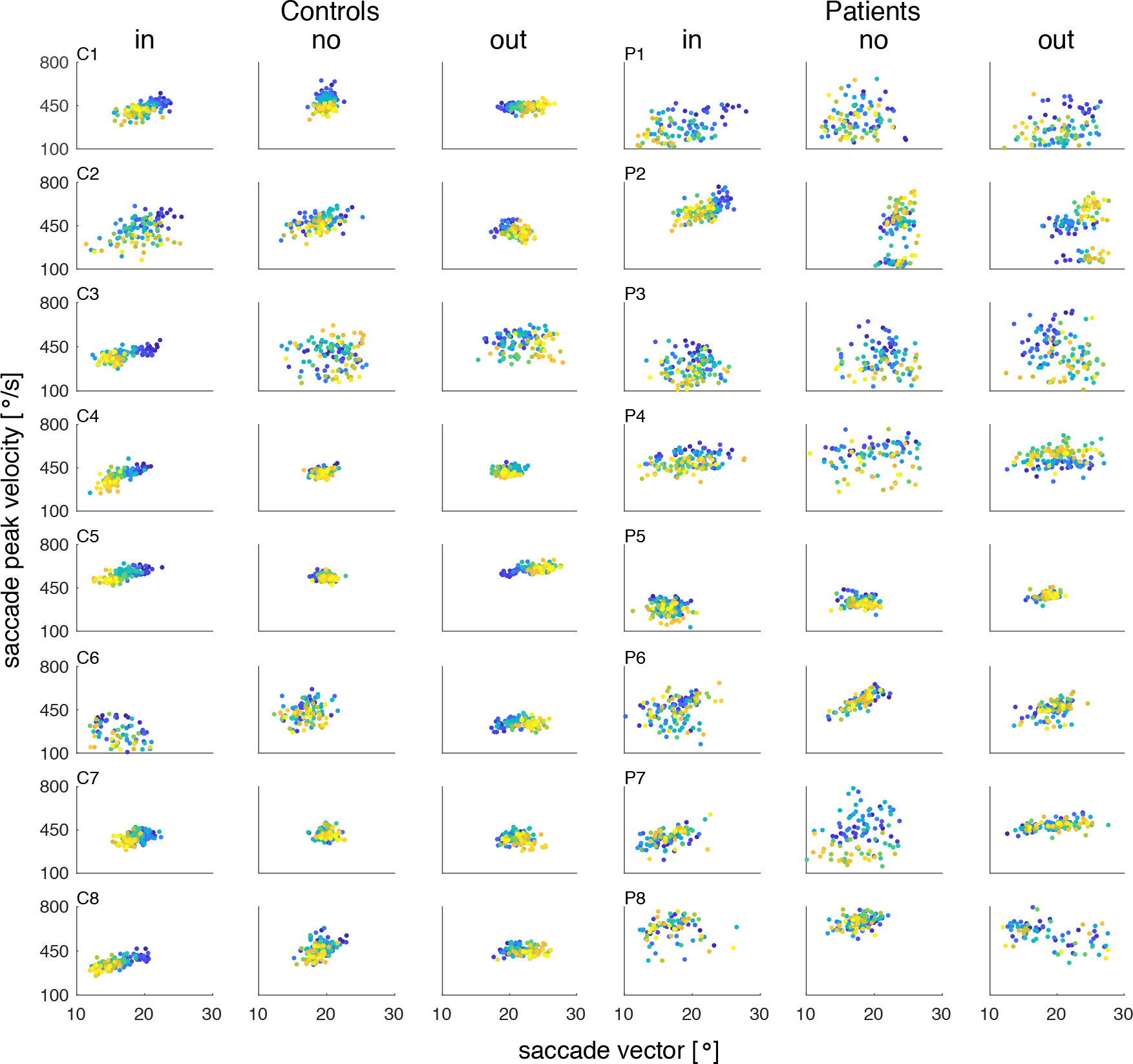
Individual saccade peak velocities in dependence on saccade vectors from early trials (blue) to late trials (yellow). Saccade peak velocities and saccade vectors are shown separately for each subject (controls C1-C8, patients P1-P8) and condition. During inward learning of control subjects, the decrease of the saccade vector encompasses a decrease of saccade peak velocity. This is in line with saccade shortening being mainly controlled by downregulating peak velocity. During outward learning, the saccade vector increases while peak velocity stays stable. This is in line with saccade lengthening being mainly controlled by upregulating saccade duration (see Fig 6). In the no step condition, saccade peak velocity encompasses a small but substantial decline that, however, is not accompanied by a decline of the saccade vector. This is a sign of oculomotor fatigue being successfully compensated by healthy subjects. As known from cerebellar patients, the patients’ saccade vectors and peak velocities are more variable than in control subjects. Moreover, overall learning effects on saccade vector and peak velocity are smaller. In the no step condition, patients show substantial oculomotor fatigue, i.e. a decline in peak velocity that is not fully compensated by saccade duration.

**Figure 6.**
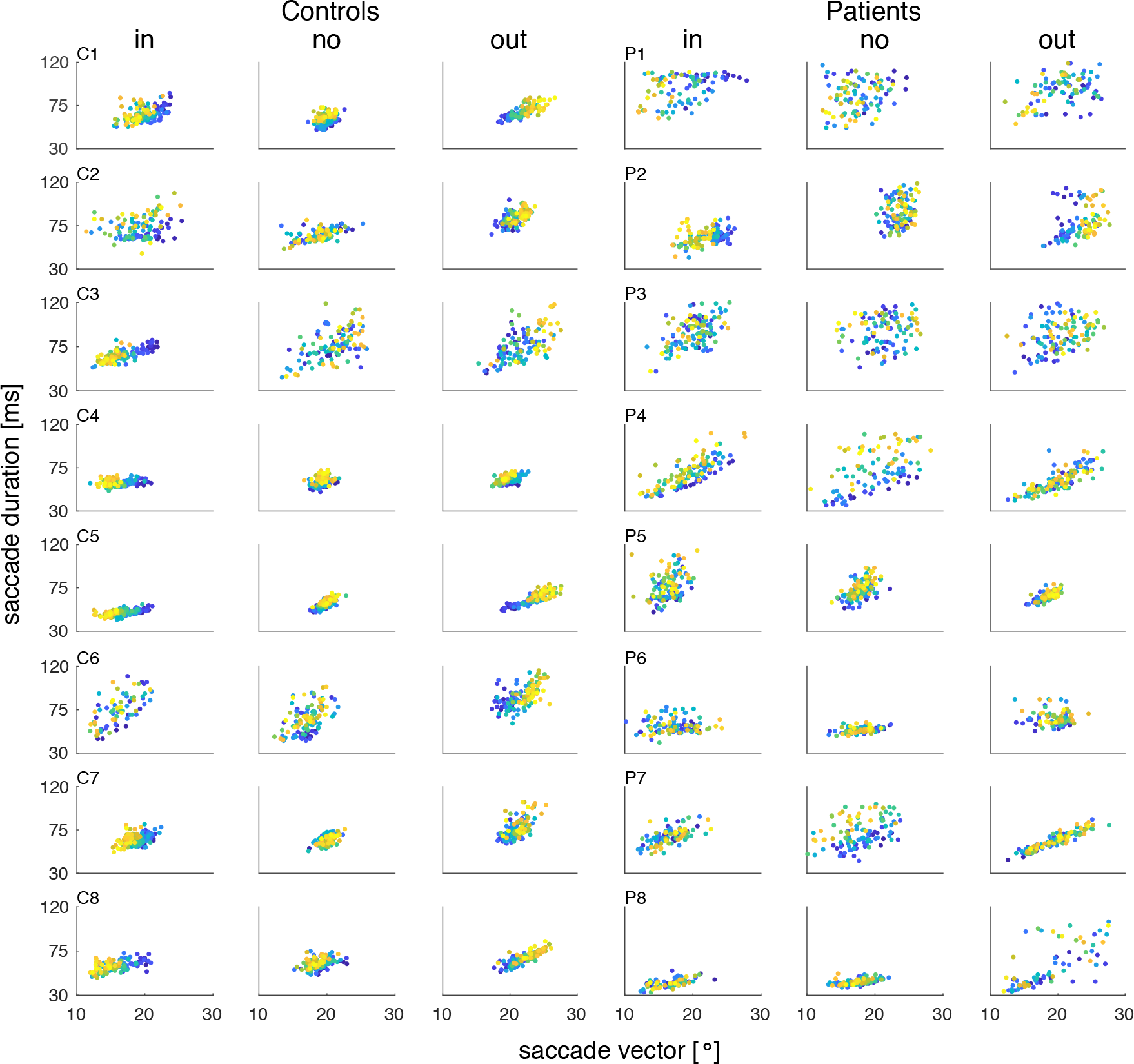
Individual saccade durations in dependence on saccade vectors from early trials (blue) to late trials (yellow). Saccade durations and saccade vectors are shown separately for each subject (controls C1-C8, patients P1-P8) and condition. During inward learning of control subjects, saccade duration declines in subjects C3, C5, C7 and C8 (yet not significantly across subjects when compared between the pre- and the post-expose phase, *t*_7_ = −1.01, *p* = .346). The decrease of the saccade vector rather stems from a decline of saccade peak velocity (see Figure 5). In the outward condition, saccade duration increases together with the saccade vector. This is consistent with saccade lengthening being mainly controlled by upregulation of saccade duration. In the no step condition, saccade duration increases, thereby compensating for the peak velocity loss seen in Figure 5. In patients, small learning effects can be seen but saccade vectors and durations are more noisy. In the no step condition, saccade duration is increased only in patient P4 and P7. Overall, duration cannot sufficiently counteract the peak velocity loss.

**Table 1.** Fitted parameters and residual standard errors. Here we report the fitted parameters and residual standard errors for the goodness of model fit, separately for each group and condition. The learning rates *α_v_*, *α_m_* and *α_cd_* were fitted to the data of the inward and the outward condition, respectively. The peak velocity decay rate *γ_k_* and the duration compensation rate *γ_λ_* were fitted to the data of the no step condition. The parameters *β_0_*, *β_k_* and *β_λ_* are the weights of the regression from saccade vectors on saccade peak velocities and saccade durations. Residual standard errors are reported separately for saccade vectors and pre- and post-saccadic localizations 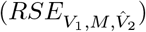, as well as for saccade peak velocities (*RSE_κ_*) and for saccade durations (*RSE_λ_*).

| Condition | Group | Fitted parameters | $\beta_0$ | $\beta_\kappa$ | $\beta_\lambda$ | $RSE_{V_1, M, \hat{V}_2}$ | $RSE_\kappa$ | $RSE_\lambda$ |
| --- | --- | --- | --- | --- | --- | --- | --- | --- |
| inward | con | $\alpha_v = 2.6*10^{-6}, \alpha_m = 2.1*10^{-5}, \alpha_{cd} = 5.9*10^{-6}$ | -9.995 | 0.034 | 0.230 | 0.8 ° | 25.0 $\frac{^\circ}{s}$ | 3.0 ms |
| | pat | $\alpha_v = 0, \alpha_m = 3.8*10^{-6}, \alpha_{cd} = 6.7*10^{-7}$ | 6.399 | 0.011 | 0.110 | 1.1 ° | 59.2 $\frac{^\circ}{s}$ | 5.1 ms |
| no step | con | $\gamma_\kappa = 0.003, \gamma_\lambda = 0.964$ | 9.828 | 0.009 | 0.088 | 0.5 ° | 23.7 $\frac{^\circ}{s}$ | 3.1 ms |
| | pat | $\gamma_\kappa = 0.008, \gamma_\lambda = 0.457$ | 8.917 | 0.009 | 0.092 | 1.1 ° | 56.1 $\frac{^\circ}{s}$ | 6.4 ms |
| outward | con | $\alpha_v = 2.3*10^{-6}, \alpha_m = 5.5*10^{-6}, \alpha_{cd} = 5.3*10^{-13}$ | -0.847 | 0.018 | 0.208 | 0.6 ° | 21.8 $\frac{^\circ}{s}$ | 3.9 ms |
| | pat | $\alpha_v = 0, \alpha_m = 1.4*10^{-6}, \alpha_{cd} = 5.1*10^{-7}$ | 8.360 | 0.006 | 0.131 | 1.1 ° | 50.8 $\frac{^\circ}{s}$ | 5.7 ms |

### Cerebellar pathology suspends short-term learning of the visuospatial target map

The fit of the visual pre-saccadic target position *V*_1_ to the pre-saccadic target localizations (Figure 3A-1,B-1, green) captures how the visuospatial target map, i.e. the visual gain *ω_v_* (Figure 3A-3,B-3, green), adapts on a short time-scale when errors are credited to an internal failure of visuospatial target representation. Consistent with previous studies (Moidell & Bedell, 1988; Collins et al., 2007; Hernandez, Levitan, Banks, & Schor, 2008; Schnier et al., 2010; Zimmermann & Lappe, 2010; Gremmler et al., 2014; Masselink & Lappe, 2021), we found a small change of *V*_1_ in learning direction for control subjects (inward −0.7 ± 2.0°, *t*_7_ = −1.00, *p* = .351; outward 1.2 ± 1.8°, *t*_7_ = 1.86, *p* = .105; Figure 3A-1). Please note that in the analysis of Cheviet et al. (2022) (which comprised more trials including those in which the flash was presented with a small distance to the 20° target position), these changes were subtly significant. Accordingly, the visual gain *ω_v_* change of control subjects was −0.04 ± 0.10 (*t*_7_ = −1.00, *p* = .351; Figure 4B) in the inward condition and 0.06 ± 0.09 (*t*_7_ = 1.86, *p* = .105; Figure 4D) in the outward condition.

By contrast, the visual pre-saccadic target position *V*_1_ of cerebellar patients did not show learning from target steps. Small changes were observed in the opposite direction (inward 0.6 ± 1.3°, *t*_7_ = 1.35, *p* = .218; outward −0.9 ± 1.1°, *t*_7_ = −2.28, *p* = .057; Figure 4B,D) but were also not significant in the analysis of Cheviet et al. (2022). Adding a mechanism that allows the model to learn in the opposite direction of error reduction seemed not justified in this context. Accordingly, we fitted a stable *V*_1_ and a stable *ω_v_* to the patients’ data (Figure 3B-1,B-3, green). In sum, patients did not show short-term learning of the visual pre-saccadic target position *V*_1_ as compared to control subjects (significant difference of *V*_1_ change between patients and control subjects in the outward condition, *t*_14_ = 2.78, *p* = .015, Figure 4D). This shows that cerebellar integrity is crucial to adapt the visuospatial target map. Neurophysiologically, the cerebellum may mediate visuospatial adaptation through feedback projections to cortical retinotopic areas (Buckner et al., 2011; Brissenden et al., 2016; Marek et al., 2018).

### Cerebellar pathology reduces short-term learning of the inverse model

The fit of the motor command *M* to the saccade vectors (Figure 3A-1,B-1, blue) quantifies how the inverse model, i.e. the motor gain *ω_m_* (Figure 3A-3,B-3, blue), adapts when errors are assigned to false computation of the motor command to bring the eyes to the visuospatial target position. In accordance with previous studies (McLaughlin, 1967; Miller, Anstis, & Templeton, 1981; Wallman & Fuchs, 1998; Bahcall & Kowler, 1999; Panouilleres et al., 2008; Ethier, Zee, & Shadmehr, 2008; Havermann & Lappe, 2010; Masselink & Lappe, 2021), the motor command *M* of control subjects decreased by −3.9 ± 1.8° during inward learning (*t*_7_ = −6.06, *p* < .001; Figure 4B) and increased by 3.6 ± 2.0° during outward learning (*t*_7_ = 5.10, *p* = .001; Figure 4D). Accordingly, the inverse model, i.e. the motor gain *ω_m_*, adapted significantly in both learning conditions (inward −0.17 ± 0.13, *t*_7_ = −3.71, *p* = .008; outward 0.12 ± 0.13, *t*_7_ = 2.5, *p* = .041). As expected by the model, the motor command changes during inward learning were transposed to saccade kinematics by downregulation of saccadic peak velocity 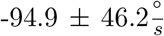, *t*_7_ = −5.81, *p* = .001; Figure 3A-5 and Figure 5) while saccade duration stayed constant (−2.6 ± 7.3 ms, *t*_7_ = −1.01, *p* = .346; Figure 3A-6 and Figure 6). By contrast, the motor command increase during outward learning was controlled by upregulation of saccade duration (10.4 ± 3.1 ms, *t*_7_ = 9.37, *p* < .001) while saccade peak velocity stayed constant 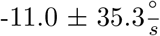, *t*_7_ = −0.89, *p* = .405). The saccade changes brought control subjects’ eyes significantly closer to the post-saccadic target (*V*_2_ change by 3.9 ± 1.8° during inward learning, *t*_7_ = 6.06, *p* < .001; −3.6 ± 2.0° during outward learning, *t*_7_ = −5.1, *p* = .001; Figure 3A-2). Thus, control subjects successfully reduced the postdictive motor error *E_post_* in both learning paradigms (inward 3.8 ± 2.4°, *t*_7_ = 4.53, *p* = .003; outward −3.3 ± 2.2°, *t*_7_ = −4.16, *p* = .004, Figure 3A-4). This corresponds to 64.1 ± 53.0% error reduction in the inward condition (*t*_6_ = 3.2, *p* = .019; Figure 4B) and 50.1 ± 36.3% error reduction in the outward condition (*t*_7_ = 3.9, *p* = .006; Figure 4D).

Compared to control subjects, motor command changes were largely reduced in cerebellar patients. During inward learning, the motor command *M* decreased by −1.2 ± 2.1° and increased during outward learning by 0.8 ± 2.6° (Figure 3B-1). These changes were not significant (inward *t*_7_ = −1.65, *p* = .143; outward *t*_7_ = 0.83, *p* = .433) and significantly less than in control subjects (inward *t*_14_ = −2.75, *p* = .016; outward *t*_14_ = 2.43, *p* = .029; Figure 4B,D). Accordingly, inverse model adaptation was not significant in patients (*ω_m_* change inward −0.08 ± 0.14, *t*_7_ = −1.73, *p* = .127, outward 0.07 ± 0.13, *t*_7_ = 1.58, *p* = .158). Saccade peak velocity was significantly decreased in the inward condition 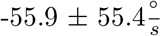, *t*_7_ = −2.85, *p* = .025; Figure 3B-5 and Figure 5) but saccade duration was not significantly increased in the outward condition (4.1 ± 13.7 ms, *t*_7_ = 0.84, *p* = .427; Figure 3B-6 and Figure 6). Hence, reduction of postdictive motor error was less effective than in the control subjects, i.e. only 5.5 ± 78.8% during inward learning (*z* = 0.84, *p* = .401; Figure 4B) and 24.5 ± 33.4% during outward learning (*t*_7_ = 2.07, *p* = .077; Figure 4D). Reduction of *E_post_* in the inward condition was significantly less in patients than in controls (*t*_14_ = 2.15, *p* = .05).

### Forward dynamics model is correctly informed about the reduced saccade change

Quantification of *CD_V_* in our model (Figure 3A-1,B-1, red) relies on the difference between pre- and post-saccadic localization, hence capturing *CD_V_*-based integration between the pre- and the post-saccadic visual scene (see Figure 3A-2,B-2 for changes of the post-saccadic target localization with respect to saccade landing, i.e. 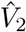. The CD gain *ω_cd_* captures how the forward dynamics model is calibrated when errors are credited to a false internal representation of the saccade.

Figure 7 shows that the motor command *M* and the *CD_V_* signal are highly correlated across control subjects and patients, respectively, in the pre-exposure phase (baseline state) as well as in the post-exposure phase of all three conditions (inward, no step, outward). This shows that the quantification of *CD_V_* based on our paradigm is a valid estimate of the internal representation of the saccade.

**Figure 7.**
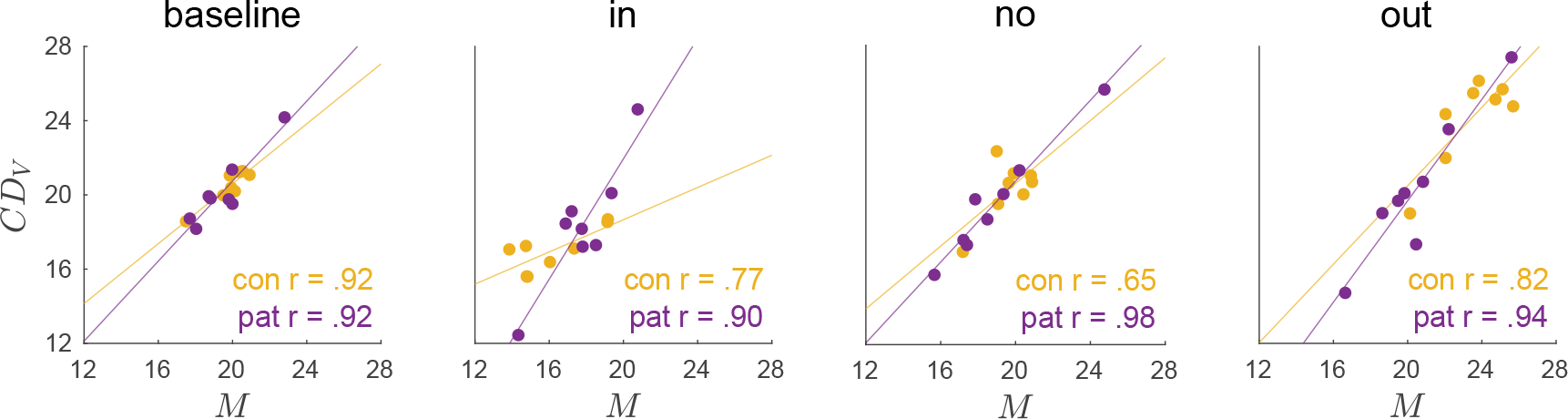
Correlation of motor command *M* and *CD_V_* signal across subjects at steady state. At steady state, i.e. in the pre-exposure phase across all three conditions (baseline) and in the post-exposure phase of the inward condition, the no step condition and the outward condition, the motor command *M* and *CD_V_* signal highly correlate across control subjects and patients. This shows that the quantification of the *CD_V_* signal by the model based on the trans-saccadic target localization and saccade data is a valid estimate of the internal representation of the saccade.

In control subjects, the *CD_V_* signal declined by −4.1 ± 2.1° during inward learning (*t*_7_ = −5.54, *p* < .001; Figure 4B) and increased by 3.9 ± 2.4° during outward learning (*z* = 2.38, *p* = .017; Figure 4D). Figure 3A-1 shows that *CD_V_* well reflects the actual saccade changes in both learning paradigms. Thus, the initial CD gain (*ω_cd_* = 1.03 ± 0.02, averaged as the baseline of all three conditions, Figure 4A) stayed stable throughout learning (inward 0.01 ± 0.08, *t*_7_ = 0.39, *p* = .707, Figure 4B; outward 0.01 ± 0.06, *z* = −0.42, *p* = .674, Figure 4D). This means that the forward dynamics model is well informed about the actual saccade changes in control subjects.

Surprisingly, this picture was similar in cerebellar patients. In both the inward and the outward condition, patients showed only a small change of the motor command *M* in learning direction that did not reach significance. This was well reflected by the *CD_V_* signal. During inward learning, *CD_V_* of patients was reduced by −1.3 ± 2.9° which did not reach significance (*t*_7_ = −1.28, *p* = .241, Figure 4B), thus well reflecting the actual saccade changes that were less than in control subjects (Figure 3B-1). In the outward condition, *CD_V_* of patients stayed roughly constant (change of −0.4 ± 3.5°, *t*_7_ = −0.35, *p* = .737, Figure 4D). Accordingly, the initial CD gain (*ω_cd_* = 1.04 ± 0.04, averaged as the baseline of all three conditions, Figure 4A) was not modified during inward learning (−0.01 ± 0.1, *t*_7_ = −0.19, *p* = .856) and only slightly modified during outward learning (−0.07 ± 0.07, *t*_7_ = −2.67, *p* = .032). In sum, the forward dynamics model seemed correctly informed about the ongoing motor changes, similar in controls and patients (no difference in *ω_cd_* change between patients and controls, inward *t*_14_ = 0.39, *p* = .704, outward *t*_14_ = 0.39, *p* = .704).

### Cerebellar pathology is accompanied by overestimation of target eccentricity

Figure 4A presents the baseline state of control subjects and patients, averaged across the three conditions. While control subjects accurately localize the pre-saccadic target on the visual map (*V*_1_ = 20.3 ± 0.8°, t-test against *P*_1_ = 20°, *t*_7_ = 1.18, *p* = .276), patients showed a significant overestimation of target eccentricity by around 1.5° (*V*_1_ = 21.5 ± 1.3°, t-test against *P*_1_ = 20°, *t*_7_ = 3.13, *p* = .017). Hence, the visual gain *ω_v_* overestimated the actual target distance already when patients entered the laboratory (*ω_v_* = 1.07 ± 0.07, t-test against 1, *t*_7_ = 3.13, *p* = .017). This was consistent across all three conditions (Figure 3B-1). Still, patients exhibited a quite accurate baseline saccade (*M* = 19.5 ± 1.6°, t-test against *P*_1_ = 20°, *t*_7_ = −0.92, *p* = .389), i.e. the motor gain *ω_m_* was downregulated such that the saccade was not hypermetric as expected from the overestimated target eccentricity (0.91 ± 0.1, t-test against 1, *t*_7_ = −2.66, *p* = .033). Interestingly, target overestimation despite a quite accurate saccade was observed before in a patient with a thalamic lesion (right posterior ventrolateral and right ventromedial, Zimmermann, Ostendorf, Ploner, & Lappe, 2015). In spite of the visuospatial hypermetria, the patients’ visuomotor system seemed to be at a steady state as the postdictive motor error *E_post_* did not significantly differ from zero (*t*_7_ = 2.11, *p* = .073 in patients, *t*_7_ = 1.73, *p* = .127 in control subjects, Figure 4A).

### Cerebellar pathology is accompanied by insufficient upregulation of saccade duration causing uncompensated oculomotor fatigue

The no step condition tested subjects’ ability to maintain a stable saccade vector when saccades were repetitively performed to the same, non-stepping target. Figure 3A-5 shows that control subjects underwent oculomotor fatigue, as revealed by the significant decline in saccade peak velocity by 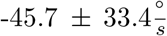 (*t*_7_ = −3.87, *p* = .006, *γ_κ_* = 0.003). However, peak velocity decline was counteracted by a significant upregulation of saccade duration (5.6 ± 5.4 ms, *t*_7_ = 2.90, *p* = .023) such that the saccade vector was maintained (*M* remained stable with *t*_7_ = 0.30, *p* = .775). The individual main sequences, i.e. the relationship of saccade peak velocity and saccade duration to the saccade vector, are shown in Figure 5 and Figure 6. With a duration compensation rate of *γ_λ_* = 0.964, saccade duration compensated for 96.4% of the within-trial velocity loss. Thus, control subjects successfully kept the visuomotor system calibrated and the error nullified (Figure 3A-4).

The cerebellar patients experienced an even greater loss of peak velocity by 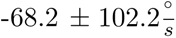 (that did not reach significance, through, probably due higher variability in patients and small sample size, *t*_7_ = −1.89, *p* = .101; *γ_κ_* = 0.008). By contrast, upregulation of saccade duration could compensate only partially (7.7 ± 14.9 ms, *t*_7_ = 1.46, *p* = .187) such that the saccade vector declined by −0.8 ± 1.5°(*t*_7_ = −1.52, *p* = .172). Even if this effect was not yet significant in the scope of only eight subjects and 120 trials, Figure 4C shows that the saccade clearly fell short in five of eight patients. Overall, saccade duration could compensate for only 45.7% (*γ_λ_* = 0.457) for the within-trial velocity loss, resulting in uncompensated oculomotor fatigue (Figure 3B-1). The deficit in saccade duration control of cerebellar patients during oculomotor fatigue appears consistent with the greater impairment during outward learning in which changes of the inverse model are usually transposed to an increase of saccade duration (Figure 3B-1 outward condition).

### Overestimation of target eccentricity may be a result of error reduction to counteract uncompensated oculomotor fatigue

We wondered how the patients’ tendency for uncompensated oculomotor fatigue affected their natural visuomotor behavior in the long run. Saccade kinematics underlie continuous fluctuations and if saccade duration cannot sufficiently compensate for regular declines of saccade peak velocity, the visuomotor system will inevitably tend to be hypometric. Saccade hypometry has been observed as a consequence of vermal cerebellar lesion in monkeys (Takagi et al., 1998; Barash et al., 1999; Ignashchenkova et al., 2009; Ohki et al., 2009) and in humans (Vahedi, Rivaud, Amarenco, & Pierrot-Deseilligny, 1995). However, Takagi et al. (1998) and Barash et al. (1999) reported that saccade vectors recovered within weeks to months. Notably, baseline saccades of cerebellar patients in our experiments were not hypometric (19.5 ± 1.6°, *t*_7_ = −0.92, *p* = .389) and not different from those of control subjects (*t*_14_ = 0.30, *p* = .767). Thus, if hypothetically, saccade hypometria occurred in the early phase of the disease, the visuomotor system would necessarily have experienced significant motor errors due to the saccade vector decline. Even if short-term error reduction is reduced in cerebellar patients, a certain ability for long-term learning is assumed to be preserved (Barash et al., 1999; Tseng, Diedrichsen, Krakauer, Shadmehr, & Bastian, 2007; Xu-Wilson, Chen-Harris, et al., 2009; Thier & Markanday, 2019). Hence, using the peak velocity decay rate and the duration compensation rate of the patients’ model fit (*γ_κ_* = 0.008, *γ_λ_* = 0.457), we simulated how the visuomotor system will behave on a long time course if, simultaneously, preserved learning counteracts the saccade decline. Figure 8 shows a simulation for 7000 saccades directed towards a 10° goal. We chose the initial visuomotor gains *ω_v_*, *ω_m_* and *ω_cd_* measured for similarly sized saccades in 35 healthy subjects by Masselink and Lappe (2021) because the larger sample size provides a more faithful picture than the initial gains of the healthy group of Cheviet et al. (2022). The simulation shows that a loss in saccade peak velocity (that is only partially compensated with *γ_λ_* = 0.457) leads to an immediate decline of the motor command *M*, hence shortening the saccade vector from 9.8° to 8.4°. In turn, the visuomotor system starts to counteract the growing motor error *E_post_*, inducing outward learning at perceptual (*ω_v_*) and motor level (*ω_m_*). Consequently, learning to counteract oculomotor fatigue leads to an outward shift of the visual pre-saccadic target localization and simultaneous recovery from saccade hypometry. The steady state at which the visuomotor system arrives is consistent with the baseline gains that we measured when patients came to the laboratory (Figure 8). In the natural situation, this time course must be longer as saccades are performed to goals of different directions and eccentricities. However, we assume that intertemporal oculomotor fatigue of multiple occurrence may lead to gradual changes of the visuomotor system similar to our simulation in the long run. In sum, the overestimation of target eccentricity in cerebellar patients may be a consequence of long-term perceptual learning to reduce motor errors.

**Figure 8.**
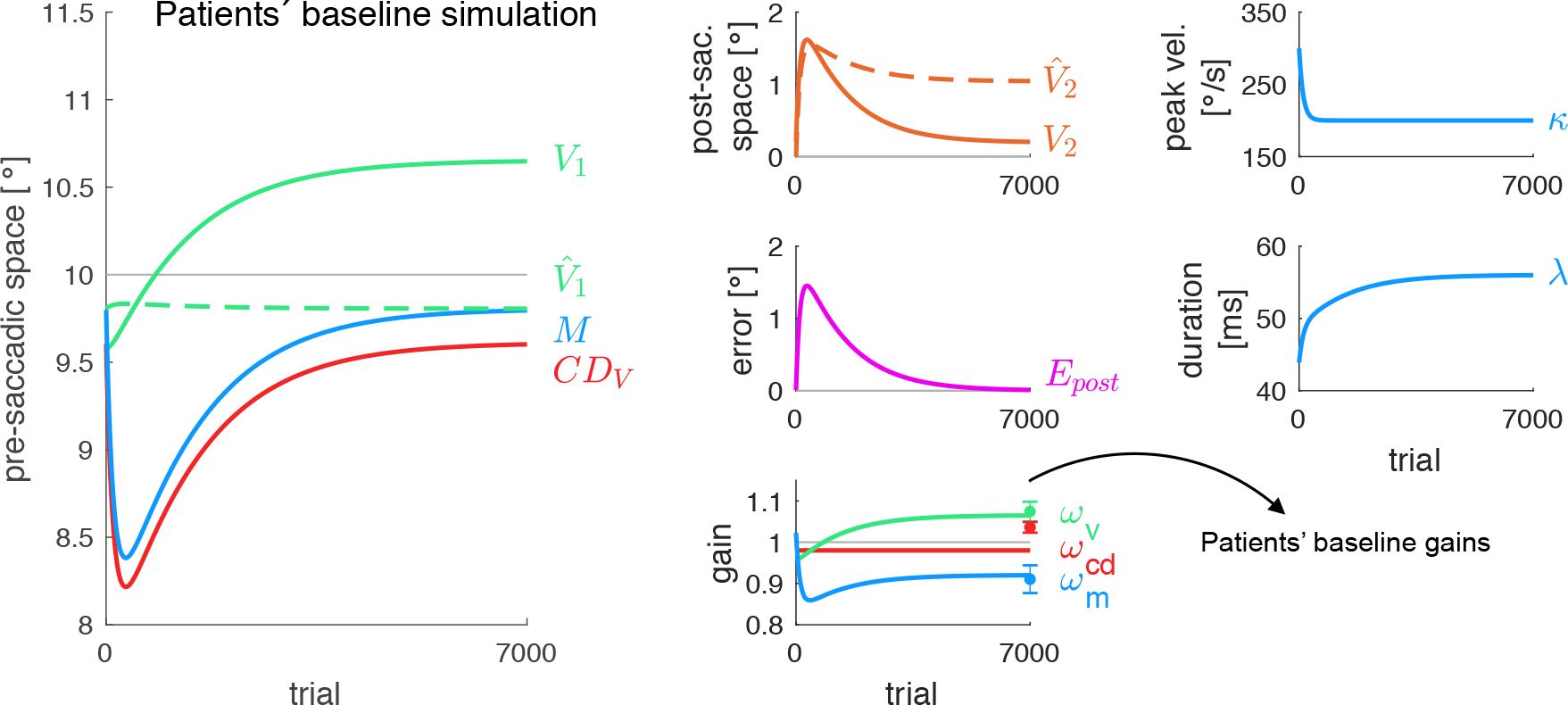
Model simulation of the patients’ baseline state. Oculomotor fatigue causes a loss of saccade peak velocity *κ* that can only be partially compensated by upregulation of saccade duration *λ*. Consequently, a decline of the saccadic motor command *M* leads to saccade hypometry, that, in turn, results in a rise of motor errors *E_post_*. Preserved long-term learning at perceptual (*ω_v_*) and motor level (*ω_m_*) counteracts to keep saccadic behavior calibrated. Hence, the visuomotor system stabilizes with an overestimation of the pre-saccadic target eccentricity (*V*_1_) and recovery from saccade hypometria (*M*), matching the baseline state measured in cerebellar patients (*ω_v_*, *ω_m_*, *ω_cd_*). Simulations were performed for a target of *P*_1_ = 10° eccentricity, ***ω***^(1)^ = (0.958; 1.023; 0.980) (which are the baseline gains measured in healthy subjects of Masselink and Lappe (2021)), 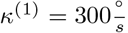, *λ*^(1)^ = 43.9 ms, *γ_κ_* = 0.008, *γ_λ_* = 0.457 (derived from the patients’ model fit to the no step condition), 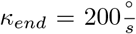, *α_v_* = 2.35*10^−6^, *α_m_* = 5.53*10^−6^, *α_cd_* = 5.31*10^−13^ (derived from control subjects’ model fit to the outward condition to simulate preserved long-term learning), *β*_0_ = −11.67, *β_κ_* = 0.03, *β_κ_* = 0.27 (derived from the saccade vectors, saccade peak velocities and saccade durations of control subjects across all three conditions to capture a minimum range of saccade vectors).

## Discussion

To dissociate cerebellar contributions to three stages of error-based learning, we modeled the visuomotor learning data of eight patients with a neurodegenerative cerebellar disease and eight healthy control subjects. The model, based on Masselink and Lappe (2021), captures recalibration (1) of the visuospatial target map, (2) of an inverse model that transforms the visuospatial movement goal into a motor command, and (3) of a forward dynamics model that estimates the saccade size from corollary discharge. The model learns from postdictive motor error, i.e. motor error with respect to a postdictive update of the pre-saccadic target position. In patients, short-term learning was reduced in the inverse model and abolished in the visuospatial target map while the forward dynamics model seemed correctly informed about the reduced saccade change. Moreover, cerebellar pathology impaired the ability to upregulate saccade duration in response to oculomotor fatigue. Modeling suggests that long-term error reduction at perceptual level may have counteracted, explaining the overestimation of target eccentricity in the patients’ baseline data. We conclude that the cerebellum recalibrates the visuospatial target map and the inverse model on a short time-scale, particularly via control of saccade duration, while the forward dynamics model does not depend on cerebellar integrity. Long-term perceptual learning may partially compensate for cerebellar motor deficits.

### Cerebellum implements short-term learning of the inverse model

Our results confirmed that the inverse model is adapted within the cerebellar microcircuit. Motor gain adaptation was largely restrained in cerebellar patients for both inward and outward errors. Similar impairments have been shown in lesioned monkeys (Takagi et al., 1998; Barash et al., 1999), in cerebellar patients (Alahyane et al., 2008; Golla et al., 2008; Xu-Wilson, Chen-Harris, et al., 2009) and in healthy subjects with cerebellar tDCS or magnetic stimulation (Panouillères, Miall, & Jenkinson, 2015; Avila et al., 2015; Jenkinson & Miall, 2010). In the patients of Golla et al. (2008), learning from outward target steps was completely suspended while in our study, a small saccade change was preserved. This difference may arise because the patients’ lesion of Golla et al. (2008) was specifically located in the oculomotor vermis, i.e. the particular region assumed to process inverse model adaptation for oculomotor commands (Hopp & Fuchs, 2004; Dean, Porrill, Ekerot, & Jörntell, 2010; Thier & Markanday, 2019).

### Cerebellum mediates short-term learning of the visuospatial target map

After visuomotor learning has initially been seen through a pure motor lens, numerous studies have revealed changes in the visuospatial perception of the movement goal (Collins et al., 2007; Schnier et al., 2010; Gremmler et al., 2014). Hence, motor learning is not simply a change of the motor command directed to the same goal location on every trial. Instead, learning underlies recalibration of both the visuospatial goal representation and the motor command required to bring the eyes to that location. This makes sense since motor errors cannot only be due to erroneous computation of the motor command but can also stem from a deficient goal representation. To our knowledge, these data are the first to demonstrate that the cerebellum is a neural substrate for visuospatial recalibration during motor learning. Neurophysiologically, the cerebellum is an ideal candidate to modulate visuospatial plasticity because of its inner circular organization (Dean et al., 2010; Tanaka, Ishikawa, Lee, & Kakei, 2020), its homogenous output organization (Sultan, Mock, & Thier, 2000; Hoshi et al., 2005; Bostan et al., 2010), and its functional interconnections with cerebral retinotopic areas (Buckner et al., 2011; Brissenden et al., 2016; Marek et al., 2018).

### The forward dynamics model is not affected by cerebellar pathology

Consistent with previous work (Bahcall & Kowler, 1999; Collins et al., 2007; Schnier et al., 2010; Masselink & Lappe, 2021), in our study, healthy subjects’ localizations shifted pre- and post-saccadically while both changes remained absent in patients. Thus, *CD_V_* seemed to correctly reflect the saccade changes in healthy controls and in patients. This result has two major implications. First, it proposes that the efference copy that is routed into the forward dynamics model reflects the actual adapted motor command and not some kind of initial motor command prior to adaptive changes. Second, it suggests that also in patients, the forward dynamics model accurately captures the actual changes of the inverse model such that the saccade vector is neither over- nor underestimated. This means that the forward dynamics model involved in trans-saccadic spatial integration and feedback motor learning remains unaffected by cerebellar degeneration. Indeed, the cerebellum computes forward models for object motion (Nowak, Hermsdörfer, Rost, Timmann, & Topka, 2004; Miall, Christensen, Cain, & Stanley, 2007; Roth, Synofzik, & Lindner, 2013) and feedforward motor control (movement adjustments during execution; Chen-Harris et al., 2008; Ethier et al., 2008; Albert & Shadmehr, 2018). However, the forward dynamics model measured in our task is one involved in trans-saccadic perception and feedback motor learning, i.e. from errors after movement completion. That this forward dynamics model may rely on areas upstream of the cerebellum is supported by studies showing that a disruption of the CD pathway from SC via MD thalamus to the FEF provokes a bias in visual space updating (Ostendorf, Liebermann, & Ploner, 2010; Prime, Vesia, & Crawford, 2010; Ostendorf, Kilias, & Ploner, 2012; Cavanaugh, Berman, Joiner, & Wurtz, 2016). The forward dynamics model could also be computed along other CD pathways, e.g. from SC via the thalamic pulvinar to parietal and occipital cortex (Wurtz, Joiner, & Berman, 2011; Berman, Cavanaugh, McAlonan, & Wurtz, 2017). The role of parietal cortex in CD processing was recently demonstrated by Cheviet, Pisella, and Pélisson (2021). At least, saccadic learning involves a distributed network across cerebral cortex (Blurton, Raabe, & Greenlee, 2012; Gerardin, Miquée, Urquizar, & Pélisson, 2012; Guillaume, Fuller, Srimal, & Curtis, 2018).

### Cerebellum counteracts oculomotor fatigue by upregulating saccade duration

Our results disclosed that cerebellar integrity is crucial for counteracting oculomotor fatigue by saccade duration adjustment. While healthy subjects compensated peak velocity loss by 96% within a trial, patients achieved only 46%, resulting in gradual saccade hypometry. This is in line with results of, firstly, Golla et al. (2008) who observed reduced oculomotor fatigue compensation, and, secondly, Xu-Wilson, Chen-Harris, et al. (2009) who observed reduced within-saccade compensation for peak velocity fluctuations in patients whose disease affected the oculomotor vermis. Taken together, it seems that saccade lengthening, within-saccade (due to oculomotor fatigue) or after saccade (due to post-saccadic motor error), is operated by cerebellar-dependent saccade duration control. That the cerebellum accommodates a velocity-duration tradeoff has been shown for hand pointing movements by Markanday, Messner, and Thier (2018). While movement duration was fine-tuned with respect to movement velocity in healthy controls, the velocity-duration tradeoff was completely absent in patients with different types of global cerebellar degeneration, i.e. comparable to our patient sample. The cerebellum adjusts movement duration in response to velocity variations also for smooth pursuit (Dash & Thier, 2013). Neurophysiologically, it remains to be examined how duration control is implemented. For manual movements, there is first evidence that the simple spikes of Purkinje cells also code movement velocity and duration (Roitman, Pasalar, Johnson, & Ebner, 2005; Pasalar, Roitman, Durfee, & Ebner, 2006; Ebner & Pasalar, 2008).

### Long-term perceptual learning may compensate for cerebellar deficiencies in saccade duration control

In the long run, insufficient velocity compensation in cerebellar patients will lead to hypometric saccades. Indeed, saccade hypometry and reduced saccade velocity were observed in cerebellar patients (Golla et al., 2008; Vahedi et al., 1995) and lesioned monkeys (Takagi et al., 1998; Barash et al., 1999; Ignashchenkova et al., 2009; Ohki et al., 2009). Even if the affected part of the cerebellum in these studies was restricted to the oculomotor vermis, we also found a hypometric tendency in the course of repetitive saccades in our patient sample. Our model simulation for long-term behavior of a cerebellar-deficient visuomotor system shows that initially, the saccade will become hypometric. Yet, with the gradual increase of motor errors, learning will counteract at perceptual and motor level. Firstly, the simulation can explain the significant overestimation of target eccentricity in the patients’ baseline data. Secondly, the simulation depicts a recovery from saccade hypometry as found in monkeys with cerebellar lesion (Takagi et al., 1998; Barash et al., 1999). Yet, in these studies, recovery concerned saccade accuracy but not saccade precision, like in our patients who exhibited non-hypometric saccades despite high endpoint variability in the baseline. Hence, a tendency for saccade dysmetria due to deficiencies in saccade duration control may induce long-term perceptual and motor learning, probably upstream of the cerebellum. This may enable the visuomotor system to stay calibrated despite cerebellar short-term adaptation deficits.

Neurophysiologically, the visual overestimation of target eccentricity is interesting when put together with Zimmermann et al. (2015) who found a similar result in a patient with a right posterior ventrolateral and ventromedial thalamic lesion. If, in both cases, this visual peculiarity is a consequence of motor adaptation deficits, these results underline the critical role of the cerebellar projection pathway via VL thalamus to frontal cortex for intact motor learning (Middleton & Strick, 2000; Gaymard, Rivaud-Péchoux, Yelnik, Pidoux, & Ploner, 2001).

### Source of saccade endpoint variability of cerebellar patients

As known from previous studies on cerebellar pathology (Takagi et al., 1998; Barash et al., 1999; Golla et al., 2008; Xu-Wilson, Chen-Harris, et al., 2009; Thier & Markanday, 2019), the cerebellar patients showed an increased variability of saccade endpoints compared to healthy controls. The source of this phenomenon is not resolved yet. Hence, the effect of increased endpoint variability on saccade motor learning remains to be examined. However, increased endpoint variability seems consistent with the role of the cerebellum in the within-saccade adjustment of movement duration in response to velocity changes that we found in the no step condition (see also Xu-Wilson, Chen-Harris, et al., 2009). If the cerebellum cannot sufficiently adjust movement duration, i.e. on the basis of feedforward saccade control (Chen-Harris et al., 2008; Ethier et al., 2008; Xu-Wilson, Chen-Harris, et al., 2009; Albert & Shadmehr, 2018), saccade endpoints will not only tend to be hypometric (in case of overall velocity decline across saccades) but will additionally be more variable as a result of trial-to-trial velocity fluctuations.

## Conclusion

We suggest an expanded view on the cerebellar circuit function for oculomotor learning and fatigue compensation. Accordingly, the cerebellum does not only adapt the inverse model at the Purkinje cell synapse, but additionally mediates the recalibration of the visuospatial goal representation on a short-time scale. The forward dynamics model for trans-saccadic space perception and feedback motor learning may be processed upstream of the cerebellum. Future research needs to examine which cerebellar structures could mediate visuospatial recalibration, and where the forward dynamics model is computed, e.g. in the thalamus, in the frontal eye fields or in other cerebral areas.

## Methods

### Subjects

Our study is based on the dataset of Cheviet et al. (2022) which includes eight healthy control subjects (49.25 ± 7.67 years, four male, four female) and eight patients with a neurodegenerative disorder of the cerebellum (55.5 ± 9.61 years, four male, four female). Three patients suffered from spinocerebellar ataxia, two patients from Friedreich ataxia, one patient from unknown autosomal dominant inherited cerebellar ataxia, one patient from fragile X–associated tremor/ataxia syndrome and one patient from late onset sporadic ataxia. Each patient was clinically examined at the Neurological Hospital Pierre Wertheimer (Bron, France) to verify a chronic progressive cerebellar ataxia, normal or corrected-to-normal vision, the ability to concentrate and to remain seated for at least 30 min as well as to maintain a stable hand position for at least 30 s. Patients with another neurological disease, unstable medical condition, psychotropic medication intake, pronounced nystagmus or ocular instability were excluded from study participation.

All subjects gave informed consent to participation. The experiment was performed in accordance with the ethical standards of the 1964 Helsinki Declaration and was approved by the ethics committee (CPP Est-III; ID-RCB: 2017–00942-51; 17.05.09).

### Setup

Subjects were seated in a completely dark room in front of a CRT monitor (19 inches, 1280*×*1024 pixels, 85 Hz, 57 cm viewing distance) that was covered with a neutral density filter to prevent visibility of monitor background light and contrast to the surrounding (ND4, 25% transmittance, 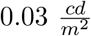 stimuli luminance, 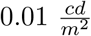 room luminance).

Movements of the right eye were recorded by an Eyelink 1000 (remote configuration; SR Research, Ontario, Canada) at 1000 Hz with a five-point calibration before each session and drift correction after each session break. Online fixation control was based on a 5° position threshold (horizontally and vertically). Saccade onset was detected online by a 2.5° position threshold, a 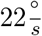 velocity threshold and a 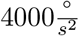 acceleration threshold. With respect to saccade initiation (0 ms), in the cerebellar group, the saccade was detected online at 16.1 ± 2.3 ms, the target was stepped (in saccade trials) or disappeared (in post-saccadic localization trials) at 28.9 ± 2.3 ms while the saccade ended at 72.7 ± 23.2 ms. In the healthy group, from saccade initiation (0 ms), the saccade was detected online at 15.5 ± 1.6 ms, the target was stepped (or disappeared) at 28.3 ± 1.5 ms while the saccade ended at 64.1 ± 3.8 ms. The procedure was controlled with Experiment Builder (SR Research, Ontario, Canada).

### Experimental design

Each subject participated in three experimental conditions recorded in randomized order with at least one week in between. Each condition measured reactive saccades to a 20° rightward target as well as visuospatial localizations of a flashed bar that varied horizontally around the target position by −4, −2, 0, +2 and +4 ° (Figure 1A). Localization was performed during fixation (pre-saccadic localization trials) and after saccade landing (post-saccadic localization trials).

The pre-exposure phase measured subjects’ baseline state, comprising four repetitions of five pre-saccadic localization trials and five post-saccadic localization trials, followed by a block of 20 saccade trials with peri-saccadic target offset (4*(5+5) + 20 = 60 trials; for an overview of the trial sequence see Figure 2A-B). The exposure phase measured 120 saccade trials with either a 6° peri-saccadic target step opposite to saccade direction (inward condition), or a 6° peri-saccadic target step in saccade direction (outward condition), or without any peri-saccadic target step (no step condition). The relatively large target step of 6° was chosen because cerebellar dysfunction is usually accompanied by a strong variability of saccade amplitudes, thereby leading to a strong variability of the post-saccadic visual target position (Takagi et al., 1998; Barash et al., 1999; Golla et al., 2008; Xu-Wilson, Chen-Harris, et al., 2009; Thier & Markanday, 2019). In order to keep this disturbance as small as possible, we chose a rather large target step, and, consequently, a rather large pre-saccadic target eccentricity of 20° such the target step was 30% of the pre-saccadic target eccentricity. Control subjects and patients reported that they did not notice the target step. In contrast to the two learning conditions, the no step condition required subjects to maintain the saccade vector, which is usually achieved by upregulating saccade duration to counteract peak velocity decline in the course of repetitive, stereotyped saccades, known as oculomotor fatigue. A break was performed after the 50th and the 100th trial.

The post-exposure phase measured localizations after exposure (four repetitions of five pre-saccadic localization trials and five post-saccadic localization trials), interleaved with three blocks of each 20 saccade refresh trials (including the respective target step) to maintain the achieved saccade vector of the exposure phase (4*(5+5) + 3*20 = 100 trials; Figure 2A-B). Each session began with a 10 min dark adaptation phase to ensure that subjects were able to discriminate the stimuli with low luminance already at the start of the session. During session breaks (before each change of trial type and after the 50th and 100th trial of the exposure phase), the room was dimly illuminated to keep the level of dark adaptation constant. The Eyelink illuminator was not visible to the participants, even after dark adaptation because the 940-nm model was used.

All stimuli (target, flash and pointer) were presented with respect to the average horizontal gaze position of a 50 ms fixation reference period at trial start (fixation reference, see trial descriptions below). This procedure ensured consistent stimulus eccentricity despite reduced fixation accuracy of cerebellar patients. A trial was repeated if fixation criteria were violated. Inter-trial interval was a 800 ms black screen.

#### Saccade trials

A red fixation circle of 1° diameter appeared for a random time interval between 800 and 1400 ms (5° to the right of the left screen border, vertically randomized between −1.5, 0 and 1.5° to the horizontal meridian, Figure 1A). If fixation criteria were met during this period, a red target circle was displayed 20° to the right with respect to the fixation reference determined during the last 50 ms of the fixation period. As soon as saccade onset was detected, the target switched off in the pre-exposure phase, and, in the exposure and post-exposure phase, stepped either 6° inward (inward condition), 6° outward (outward condition) or stayed at its position (no step condition). In the exposure and post-exposure phase, the target stayed visible for 100 ms.

#### Post-saccadic localization trials

A red fixation circle of 1° diameter was presented 5° to the right of the left screen border on the horizontal meridian (Figure 1A). Subjects were instructed to fixate the circle and to indicate when ready to begin the trial via mouse press. If, after a 450 ms black screen, fixation criteria were met for 50 ms, a blue vertical line was flashed with counterbalanced horizontal distance of 16, 18, 20, 22 or 24° with respect to the fixation reference determined during the 50 ms fixation check period (12 ms, 0.2° width, 29° height). After a 400 ms black screen, a red target circle of 1° diameter appeared 20° to the right of the fixation reference. As soon as saccade onset was detected, the target was extinguished, followed by a 100 ms black screen. A blue line pointer was displayed on the right screen bottom (0.2° width, 10° height, 10° below the horizontal meridian, randomly 17, 19, 21 or 23° to the right of the fixation reference). As soon as subjects started moving the mouse, the pointer was blocked to the horizontal meridian. The subject had to move the pointer to the perceived horizontal flash position, confirming the judgement with a mouse press.

#### Pre-saccadic localization trials

A red prohibited direction sign served as a fixation stimulus (with the same spatial parameters as in the post-saccadic localization trials, Figure 1A). The subject had to continuously fixate it from the first mouse press (to indicate she/he is ready to start the trial) to the second mouse press (to confirm the localization judgement). The flash was presented with the same latency and spatial parameters as in the post-saccadic localization trials. The pointer appeared 750 ms after flash offset to roughly match the respective interval of the post-saccadic localization trials, and with the same latency and spatial parameters as in the post-saccadic localization trials. As soon as the subject started moving the mouse, the pointer was blocked to the horizontal meridian and the subject had to move the pointer to the perceived horizontal flash position and press the mouse.

#### Training phase

Before each session, subjects practiced a minimum of 5 pre- and 5 post-saccadic localization trials. In case of difficulties, online gaze position and a fixation box around the fixation circle were displayed to help subjects to familiarize with the task requirements.

### Model

To quantify the adaptive changes in the visuomotor circuitry of patients and control subjects, we used the model of Masselink and Lappe (2021) and expanded it with (1) how changes in the motor command are transposed to saccade peak velocity and saccade duration and (2) how the visuomotor system compensates for a fatigue-induced decline in saccade peak velocity (Figure 1B). The model describes the saccadic circuitry based on three sensorimotor transformations, i.e. synaptic gains (Shadmehr et al., 2010; Wolpert et al., 2011), that learn to reduce postdictive motor error, an error of the motor command with respect to a postdictive update of the pre-saccadic target position. These plastic gains are, first, a visual gain *ω_v_* to transform retinal input into target position on a visuospatial map, second, a motor gain *ω_m_* to transform spatial target position into a motor command (inverse model), and, third, a CD gain *ω_cd_* to transform the corollary discharge *CD_M_* of the motor command into *CD_V_*, i.e. the computed displacement of visual space due to the saccade (forward dynamics model).

#### Pre-saccadic calculations

The pre-saccadic target position *V*_1_^(*n*)^ on the visuospatial map is scaled by the visual gain *ω_v_*^(*n*)^:

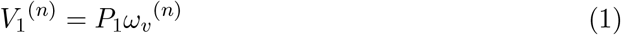

with the trial number *n* and physical target eccentricity *P*_1_ (Figure 1B-1). If *ω_v_*^(*n*)^ = 1, the target is accurately localized on the visuospatial map. If a postdictive motor error is assigned to an internal failure of visuospatial target representation, *ω_v_*^(*n*)^ is adapted to reduce the error for future movements (see equation 14).

The inverse model (Figure 1B-2) transforms the pre-saccadic target position into a motor command with the motor gain *ω_m_*^(*n*)^:

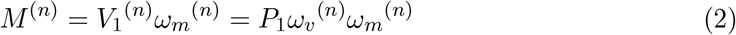

Thus, the inverse model transforms the movement goal from visuospatial to motor coordinates based on the assumed state of muscle dynamics. If *ω_m_*^(*n*)^ = 1, the saccade lands at the spatial position *V*_1_^(*n*)^. If a postdictive motor error is assigned to a change in muscle dynamics, *ω_m_*^(*n*)^ is adapted to reduce the error for future movements (see equation 14). The quantification of changes to the visual gain *ω_v_*, on the one hand, and the motor gain *ω_m_*, on the other hand, dissociates the contribution of the two processes to behavioral output, i.e. saccade vector change.

The motor command *M* ^(*n*)^ is copied into *CD_M_* ^(*n*)^:

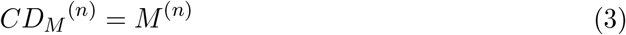

and directed into the CD pathway.

Before saccade start, the forward dynamics model (Figure 1B-3) transforms the corollary discharge of the motor command *CD_M_* ^(*n*)^ into the *CD_V_* ^(*n*)^ signal, i.e. the computed displacement of visual space, to inform the visual system about the upcoming saccade:

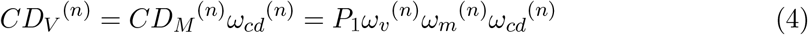

If *ω_cd_*^(*n*)^ = 1, *CD_V_* ^(*n*)^ matches the actual saccade vector. If a postdictive motor error is assigned to an internal failure of *CD_V_* representation, *ω_cd_*^(*n*)^ is adapted to reduce the error for future movements (see equation 14).

Based on *CD_V_* ^(*n*)^, the forward outcome model (Figure 1B-4) shifts the coordinates of the visual pre-saccadic target position *V*_1_^(*n*)^ to predict the visual post-saccadic target position:

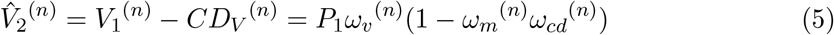

#### Saccade kinematics

Saccadic motor learning has been shown to be differently transposed to saccade kinematics, i.e. to saccade peak velocity and saccade duration, depending on the error direction (Golla et al., 2008; Thier & Markanday, 2019). Moreover, there is evidence that cerebellar patients are more impaired in learning from outward than from inward errors (Golla et al., 2008). Thus, capturing the learning effect on the patients’ saccade kinematics may provide a deeper understanding of how the cerebellum operates the recalibration of motor commands. Hence, we expanded the model of Masselink and Lappe (2021) with how the motor command is transposed to saccade peak velocity and saccade duration. Accordingly, the saccade is executed with the saccade peak velocity *κ*^(*n*)^ and the saccade duration *λ*^(*n*)^ that are specific for the motor command (Figure 1B-5, see equations 16, 18 and 20 for how the motor command is transposed to saccade peak velocity and saccade duration depending on the paradigm). This specificity relies on a linear relationship between a saccade’s amplitude, its peak velocity and its duration, known as the saccadic main sequence (Bahill et al., 1975; Harris & Wolpert, 2006; Collins, Semroud, Orriols, & Doré-Mazars, 2008). Hence, we specify the relationship between the motor command *M* ^(*n*)^, the saccade peak velocity *κ*^(*n*)^ and the saccade duration *λ*^(*n*)^ by the following equation:

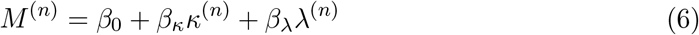

For the modeling, we derive *β*_0_, *β_κ_* and *β_λ_* from fitting the equation to the saccade vectors, saccade peak velocities and saccade durations of patients and healthy controls. Please note that we use one single function to express the relationship of saccade amplitude, peak velocity and duration in order to be able to model oculomotor fatigue compensation, i.e. when a decrease of saccade peak velocity does not inevitably result in a decrease of the saccade amplitude but can be compensated by a longer saccade duration such that the saccade amplitude stays stable. We used a linear approximation in sort of a 3D plane as described by equation 6 that appeared well justified over the range of saccade amplitudes that we measured. Two separate functions expressing the relationship of the saccade amplitude, first, to peak velocity and, second, to duration (as e.g. used by Bahill et al., 1975; Becker, 1989) would not allow to model oculomotor fatigue compensation.

#### Saccade execution

The execution of the motor command produces the physical saccade vector (Figure 1B-6):

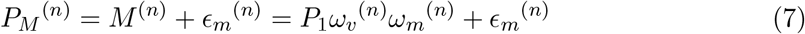

with random motor noise *ϵ_m_*^(*n*)^.

In the inward and outward learning paradigms, the target is shifted during saccade execution. The peri-saccadic target displacement due to the target shift *P_s_* and the motor execution noise *ϵ_m_* is described by:

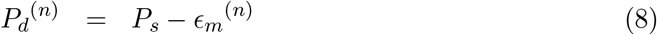

#### Post-saccadic calculations

After saccade offset, the post-saccadic target is localized on the visuospatial map (Figure 1B-7):

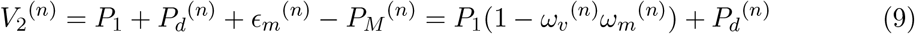

Please note that the visual gain *ω_v_*^(*n*)^ is not applied to *V*_2_^(*n*)^ because visuospatial changes during saccadic motor learning underlie an adaptation field (Collins et al., 2007; Schnier et al., 2010). Hence, a change in the visual gain *ω_v_*^(*n*)^ is a local effect around the pre-saccadic target position and should not be applied to the representation of the post-saccadic target after saccade landing.

The backward outcome model integrates the visual post-saccadic target position *V*_2_^(*n*)^ with *CD_V_* ^(*n*)^ to from a postdictive update of the target position in a pre-saccadic reference frame (Figure 1B-8):

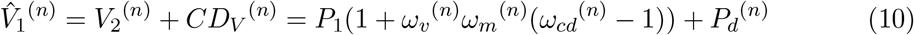

We refer to this process as the backward outcome model as it performs a postdictive transformation from post- to pre-saccadic space, analogously to the forward outcome model that performs a predictive transformation, i.e. from pre- to post-saccadic space.

#### Learning of visuomotor gains and its transposition to saccade kinematics

In the inward and outward learning paradigms, the visuomotor system experiences a significant motor error due to the peri-saccadic target step. This error, termed the postdictive motor error *E_post_*^(*n*)^, is calculated as the error of the motor command with respect to the postdicted pre-saccadic target position (Figure 1A-9):

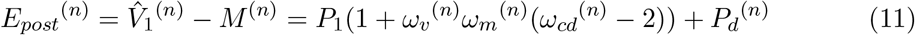

To reduce *E_post_*^(*n*)^ on the next trial, the visuomotor system adapts its gains following an internal estimate of the *E_post_*^(*n*)^ gradient (Doya, 1999; Wolpert et al., 2011; Taylor & Ivry, 2014). We express this principle by the delta rule (Widrow & Hoff, 1960; Widrow & Stearns, 1985):

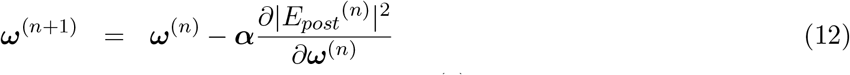

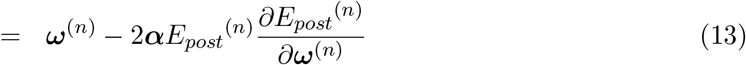

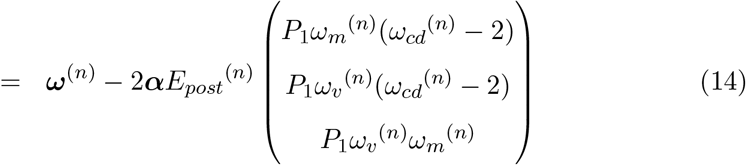

The learning rates 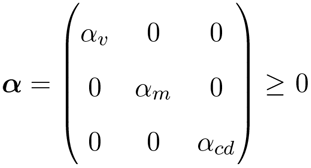 specify the speed of learning.

The changes in the visual gain *ω_v_* and in the motor gain *ω_m_* result in a changed motor command *M* ^(*n*+1)^ on the next trial. The transposition of the changed motor command to saccade peak velocity and saccade duration needs to take into account whether the motor command was decreased or increased, i.e. whether changes occurred during inward or during outward learning (Golla et al., 2008; Thier & Markanday, 2019). Saccade shortening during inward learning is usually characterized by a decline of peak velocity while saccade duration stays constant. Accordingly, during inward learning in our model (*P_s_* < 0), saccade duration *λ* stays constant while saccade peak velocity *κ* is downregulated on the next trial according to the changed motor command (Figure 1B-6):

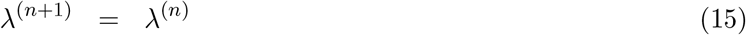

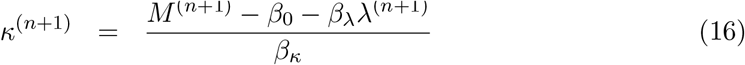

By contrast, saccade lengthening during outward learning is usually characterized by an increase of saccade duration while saccade peak velocity stays constant (Golla et al., 2008; Thier & Markanday, 2019). Accordingly, during outward learning in our model (*P_s_* > 0), saccade peak velocity *κ* stays constant while saccade duration *λ* is upregulated on the next trial according to the changed motor command (Figure 1B-6):

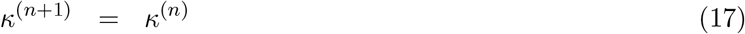

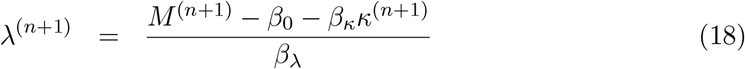

The visuomotor system reaches a steady state, i.e. learning comes to an end, if the saccade lands at the postdicted pre-saccadic target position such that *E_post_*^(*n*)^ = 0.

#### Oculomotor fatigue and its compensation by saccade duration

For a deeper understanding of which calibration processes are perturbed due to cerebellar disease, we also modeled the experimental condition without any target step, i.e. without learning. Although target localizations and saccade vectors should stay roughly constant in healthy subjects, the repetitive execution of saccades to the same target position is usually accompanied by a decline in saccade peak velocity, a sign of oculomotor fatigue. However, this compensating mechanism has been shown to be impaired in patients with vermal cerebellar lesion (Golla et al., 2008). Accordingly, in the no step condition of our model (*P_s_* = 0), saccade peak velocity *κ* declines while saccade duration *λ* compensates by a certain percentage on the next trial (Figure 1B-6):

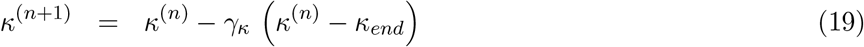

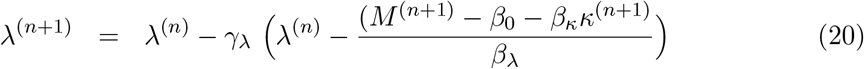

Here, *γ_κ_ ≥* 0 describes the decay rate of peak velocity, *κ_end_* limits peak velocity loss and *γ_λ_* describes the percentage by which saccade duration compensates for the peak velocity loss (0 *≤ γ_λ_ ≤* 1). According to the changes of peak velocity *κ* and duration *λ* in the no step condition (*P_s_* = 0), the motor gain *ω_m_*^(*n*+1)^ of the next trial is:

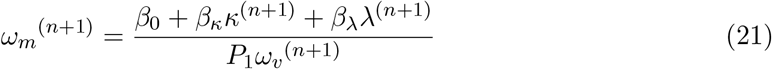

Hence, during expected oculomotor fatigue in the no step condition, we model changes in saccade peak velocity and saccade duration that are reflected in the motor gain *ω_m_* while *ω_v_* and *ω_cd_* are assumed to stay stable.

### Analysis

#### Data processing

Data analysis was performed in Matlab R2017a (Mathworks, Natick, MA). Saccades were detected offline with a self-made routine that searched for a saccade after target onset with an onset and offset velocity threshold of 80 deg/s. Saccade traces as well as onset and offset detection were visualized trial by trial in order to correct for detection errors. Based on the verified saccade onset and offset, saccade latency, peak velocity and duration were extracted. The horizontal eye position signal was smoothed and the saccade start position was determined based on the position 50 ms prior to saccade onset detection. The saccade end position was extracted based on the position 50 ms after saccade offset detection to prevent post-saccadic oscillation effects.

Saccades with a latency of less than 100 ms, an amplitude below 10° or above 30° as well as localizations with more than ±7° localization error were excluded from the analysis. Moreover, as our modeling approach explicitly relies on the localization judgement of target eccentricity, we chose only the pre- and post-saccadic localization trials in which the flash position matched the saccade target position, i.e. 20° eccentricity. Thus, we ensured that our modeling approach quantifies perceptual changes exactly at the spatial position of the target and is not disturbed by the process of learning transfer to nearby locations, known as the saccadic adaptation field (Deubel, 1987; Miller et al., 1981; Frens & Van Opstal, 1994; Straube et al., 1997; Hopp & Fuchs, 2004; Collins et al., 2007; Schnier et al., 2010; Pelisson et al., 2010).

For each condition of each subject, we calculated the mean saccade vector, mean saccade peak velocity and mean saccade duration of the 20 pre-exposure saccade trials and the last 20 exposure saccade trials, respectively. For the pre- and the post-saccadic flash localizations we extracted the mean of the pre-exposure and the post-exposure phase. In addition, we quantified a cross-condition baseline state for each subject, averaging the mean saccade vectors, saccade peak velocities, saccade durations and pre- and post-saccadic localizations across the three conditions. Outliers with more than three standard deviations from the mean were excluded. We averaged the saccade vectors, pre- and post-saccadic localizations at group level, i.e. separately for patients and control subjects and condition. Moreover, we calculated the mean trial-by-trial saccade vectors, saccade peak velocities and saccade durations of the 120 exposure trials, separately for patients and control subjects.

#### Model-based analysis

We calculated the state of the *CD_V_* signal, the visuomotor gains ***ω*** and the *E_post_* error of each subject’s pre-exposure and post-exposure phase. Based on the pre-saccadic target localization (*V*_1_), the saccade vector (*M*) and the post-saccadic target localization with respect to the saccade landing position 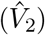, we derive:

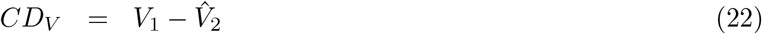

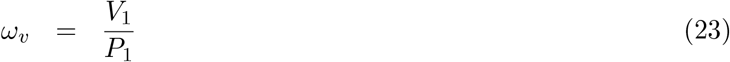

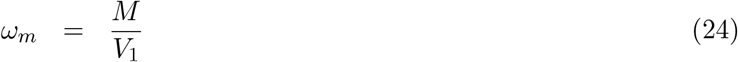

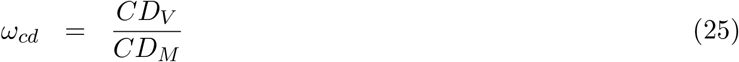

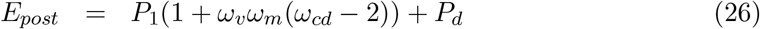

with *P*_1_ = 20° and *P_d_* = 0° for the pre-exposure phase and the post-exposure phase of the no step condition, *P_d_* = −6° for the post-exposure phase of the inward condition and *P_d_* = 6° for the post-exposure phase of the outward condition. Additionally, we derived the change of each variable from the pre- to the post-exposure phase.

Before fitting the model to the trial-by-trial data, we performed a regression of the saccade vectors on saccade peak velocities and saccade durations of the exposure phase, separately for each group (controls, patients) and each condition (inward, no step, outward). See Table 1 for the fitted regression weights.

Starting from the pre-exposure mean of pre-saccadic localization (*V*_1_^(1)^), post-saccadic localization (with respect to the saccade landing point, 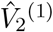, saccade vector (*M* ^(1)^), saccade peak velocity (*κ*^(1)^) and saccade duration (*λ*^(1)^) taken as values of trial *n* = 1, we fitted the model to the exposure phase, separately for each group (control subjects, patients) and each condition (inward, no step, outward). For the inward and the outward condition, we fitted the learning rates of the visuomotor gains (***ϕ_fit_*** = ***α***), and for the no step condition, we fitted the peak velocity decay rate and the duration compensation rate (***ϕ_fit_*** = (*γ_κ_*, *γ_λ_*)). The fitting procedure minimized the weighted sum of squared errors (SSE):

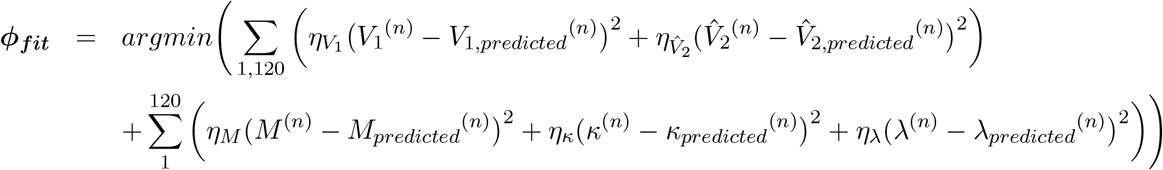

with the weights 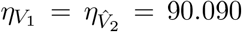, *η_M_* = 0.015, *η_κ_* = 0.012 and *η_λ_* = 0.003 to account for the unequal number of data points the signals were fitted to (e.g. only two data points for *V*_1_ and 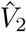 but 120 data points for *M*, *κ* and *λ*) and for the unequal units (e.g. a saccade vector error of 3° should be weighted more than a peak velocity error of 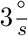. In the inward and in the outward condition patients showed a small *V*_1_ change in the opposite direction of learning. This effect was small, not significant and, thus, rather of random nature (see Figure 3B). In these two cases, we fitted a stable *V*_1_ as the mean of the pre-saccadic target localizations between the pre- and the post-exposure phase. For all model fits, we set *ϵ_m_*^(*n*)^ = 0.

For goodness of fit, residual standard errors *RSE* were calculated, collectively for *V*_1_, *M* and 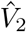 (in °) and separately for 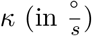 and *λ* (in ms):

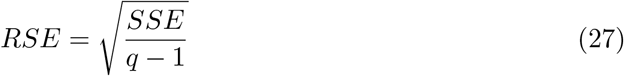

with *q* as the number of data points used for the respective *RSE* calculation.

#### Statistical analysis

To compare the data of one group against zero, we used two-sided one-sample t-tests or a Wilcoxon signed rank test in case normality distribution was violated. To compare data between groups, i.e. between patients and control subjects, two-sided two-sample t-tests, or, alternatively, Wilcoxon rank sum tests in case normality distribution was violated, were calculated. All tests were conducted with a significance level of 0.05.

## Acknowledgements

We thank Pr. Caroline Froment-Tilikete for helping us in the recruitment and clinical evaluation of the patients, and Eric Koun and Roméo Salemme for their assistance with the technical implementation of the protocol and their advice about eye movements analysis tools.

